# Antioxidant nanozyme counteracts HIV-1 by modulating intracellular redox potential

**DOI:** 10.1101/2020.08.15.252098

**Authors:** Shalini Singh, Sourav Ghosh, Virender Kumar Pal, MohamedHusen Munshi, Raju S Rajmani, Govindasamy Mugesh, Amit Singh

## Abstract

Reactive oxygen species (ROS) regulates the proliferation of human immunodeficiency virus (HIV-1) and *Mycobacterium tuberculosis* (*Mtb*) inside the infected immune cells. However, the application of this knowledge to develop therapeutic strategies remained unsuccessful due to unfavorable consequences of manipulating cellular antioxidant systems that respond to ROS. Here, we show that vanadium pentoxide (V_2_O_5_) nanosheets functionally mimic the activity of natural glutathione peroxidase (GPX) to mitigate ROS associated with HIV-1 infection without triggering detrimental changes in cellular physiology. Using genetic reporters of glutathione (GSH) redox potential (*E_GSH_*; Grx1-roGFP2) and H_2_O_2_ (Orp1-roGFP2), we showed that V_2_O_5_-nanosheets catalyze GSH-dependent neutralization of ROS in HIV-1 infected cells. Notably, V_2_O_5_-nanosheets uniformly blocked HIV-1 reactivation, multiplication, and impaired survival of drug-resistant *Mtb* during HIV-TB co-infection. Mechanistically, V_2_O_5_-nanosheets suppressed HIV-1 by affecting the expression of pathways coordinating redox balance, virus transactivation (*e.g.,* NF-κB and FOS), inflammation, and apoptosis. Importantly, a combination of V_2_O_5_-nanosheets with a pharmacological inhibitor of NF-κB (BAY11-7082) abrogated activation of HIV-1 from latency. Lastly, V_2_O_5_-nanosheets counteracted ROS, disease pathophysiology, and virus expression in HIV-1 transgenic mice. Our data successfully revealed the usefulness of V_2_O_5_-nanosheets against human pathogens and suggest nanozymes as future platforms to develop interventions against infectious diseases.

**Significance Statement:** Redox stress, such as those caused by the deregulation of the antioxidant glutathione, promotes the multiplication of human immunodeficiency virus-1 (HIV-1) and *Mycobacterium tuberculosis* (*Mtb*). Here we present a vanadium pentoxide (V_2_O_5_)-based antioxidant nanozyme that targets cells infected with HIV-1. The nanozyme, by mimicking the activity of glutathione peroxidase, reprograms redox signaling to subvert HIV-1 from monocytes, lymphocytes, and HIV-1 transgenic mice. Treatment with nanozyme bolsters the antiviral potential of immune cells by reducing the expression of genes involved in virus activation, inflammation, and apoptosis. The nanozyme also inhibited the proliferation of *Mtb*, which is a major cause of lethality in HIV patients. These V_2_O_5_-based nanozymes may be applied to numerous human pathogens where redox signaling contributes to disease progression.

## Introduction

Nanomaterials with intrinsic enzyme-mimetic properties (nanozymes) have been explored as low-cost alternatives to natural enzymes (*1*). The application of these nanomaterials was largely restricted to industries for chemical synthesis, detection of biomolecules, and bioremediation (*1, 2*). While largely ignored by the biomedical community, recent studies provide evidence for the clinical importance of artificial nanozymes *in vivo* (*3*). For example, ceria-based nanoparticles (NPs) mimic superoxide dismutase (SOD) activity and exhibit neuroprotection and reduced inflammation (*4, 5*). Similarly, iron oxide-based nanoparticles mimic peroxidase–like activity and protect from bacterial biofilms associated with oral infection (*6*). Moreover, ferumoxytol, an FDA approved iron-oxide nanoparticle has been shown to inhibit tumor growth in mice (*7*). Recently, we reported that vanadium pentoxide (V_2_O_5_) nanomaterial could protect mammalian cells from oxidative damage and apoptosis by glutathione peroxidase (GPX)-like activity in the presence of cellular antioxidant glutathione (GSH) (*8*). These findings indicate the importance of nanozymes mimicking the activity of natural antioxidant enzymes for biomedical applications.

Our particular interest is to apply antioxidant nanozymes in oxidative stress linked human infections where therapeutic options are inadequate. In this context, oxidative stress is central to the infection caused by one of the major global pathogen human immuno-deficiency virus (HIV-1; the causative agent of the acquired immuno-deficiency syndrome [AIDS]) (*9*). A major barrier to curing HIV-1 infection is latency, wherein the infected cells harbor the intact viral genome that is replication-competent but transcriptionally silent. Interestingly, oxidative stress is known to reactivate HIV-1 from latent reservoirs via NF-κB directed transcriptional activation of the viral long terminal repeat (LTR) (*10, 11*). Further studies linking oxidative stress with HIV-1 infection demonstrate variations in GSH levels in infected cells and tissues (*12–14*). Using a non-invasive biosensor of GSH redox potential (*E_GSH_*_;_ Grx1-roGFP2), we discovered that reductive *E_GSH_* sustains viral latency, whereas a marginal oxidative shift in *E_GSH_* promotes HIV-1 reactivation (*15*).

In addition to GSH, decreased expression levels of host GPXs are associated with HIV-1 reactivation and replication (*15, 16*). The HIV-1 genome also encodes a fully functional GPX (HIV-1 vGPX) module (*17*), which protects cells from reactive oxygen species (ROS) induced apoptosis and possibly helps HIV-1 to maintain latency (*17*). Consistent with this, majority of the viral isolates from individuals naturally competent to maintain HIV-1 in a latent state (long-term non-progressors [LTNPs]) possess a functional vGPX gene (*18*). In contrast, HIV-1 strains from the patients with actively replicating virus contain a non-functional vGPX (*18*). Since LTNPs serve as a natural model for slowing progression of HIV in humans, the association of functional vGPX with these individuals reinforces the physiological importance of the GSH/GPX axis in HIV-1 latency. Despite these observations, efforts to mitigate oxidative stress for subverting HIV-1 reactivation by either supplementation of GSH precursor (N-acetylcysteine [NAC]) or activation/over-expression of GPXs yielded inconsistent results (*19–22*). While counterintuitive, these findings are in agreement with several studies showing the adverse influence of uncontrolled overproduction of natural antioxidant systems (including GPXs) on redox metabolism and disease outcome (*22*). Further, over-expression of GPXs might not be sufficient as cellular GPX activity is enhanced by oxidative stress responsive post-translational modifications (PTMs) such as phosphorylation, carbonylation and O-GlcNAcylation (*23–25*). Besides this, the activity of GPXs is also dependent on the availability of selenium (Se), an essential micronutrient that has been reported to be low in HIV patients (*16*). In this context, artificial nanozymes mimicking GPX-like activity (*e.g.,* V_2_O_5_ NPs) with high sensitivity and specificity under physiological conditions found in the human body (*i.e.* mild temperature, pH 4- 8, and aqueous buffer) can provide a suitable alternate to natural GPXs. We envisage that antioxidant nanozymes can be exploited to generate new knowledge on redox signaling mechanisms underlying HIV-1 latency and reactivation, which could aid the development of fresh therapeutic approaches for targeting HIV.

In this work, we exploited multiple interdisciplinary approaches to describe the utility of V_2_O_5_ thin-nanosheet (Vs) displaying functional GPX activity in dissecting redox signaling underlying HIV-1 latency. By exploiting genetic reporters, cellular models of HIV-1 latency, and HIV transgenic mice (Tg26), we discovered that Vs- mediated remediation of ROS efficiently subverted reactivation of HIV-1 and survival of another human pathogen (*Mycobacterium tuberculosis* [*Mtb*]) that frequently infects HIV patients. Our study provides an elegant example of how antioxidant nanozymes can be explored to understand the redox basis of human infections.

## Results

### Synthesis and biophysical characterization of catalytically efficient V_2_O_5_ thin-nanosheets (Vs)

We have recently shown that the V_2_O_5_ nanomaterials exhibit excellent isoform-specific GPX activity, which is dependent on the surface exposed crystal facets (*26*). To make V_2_O_5_ nanomaterials biologically useful, we focused on two distinct morphologies of V_2_O_5_ nanomaterials i.e. wires (VNw) and sheets (VSh) (Fig. S1, A and B). Both the morphologies displayed similar surface-exposed crystal facet {001}. However, VSh exhibited a higher GPX reaction rate as compared to VNw as determined by measuring the decrease in NADPH absorbance at 340 nm using glutathione reductase (GR)-coupled assay (see *Materials and Methods*) (Fig. S1C). Smaller nanoparticles often display higher catalytic activity (*27–31*). Therefore, we made subtle modifications in VSh preparation to generate ultrathin-nanosheets (Vs) (Fig. 1A). We confirmed the morphology of Vs by scanning electron microscopy (SEM) (Fig. 1B). Examination of the rate of GPX activity revealed that Vs possesses a 1.6- to 2-fold higher capacity to reduce H_2_O_2_ in the GR-coupled assay (Fig. 1C) as compared to VNw and VSh (Fig. 1D). A comparison of activities with three different peroxides - H_2_O_2_, tertiary-butyl hydroperoxide (*t-*BuOOH) and cumene hydroperoxide (Cum-OOH), indicates that Vs is very selective towards H_2_O_2_ (Fig. 1E). Based on this, we carried out the extensive biophysical and biochemical characterization of Vs *in vitro*.

**Figure 1.**
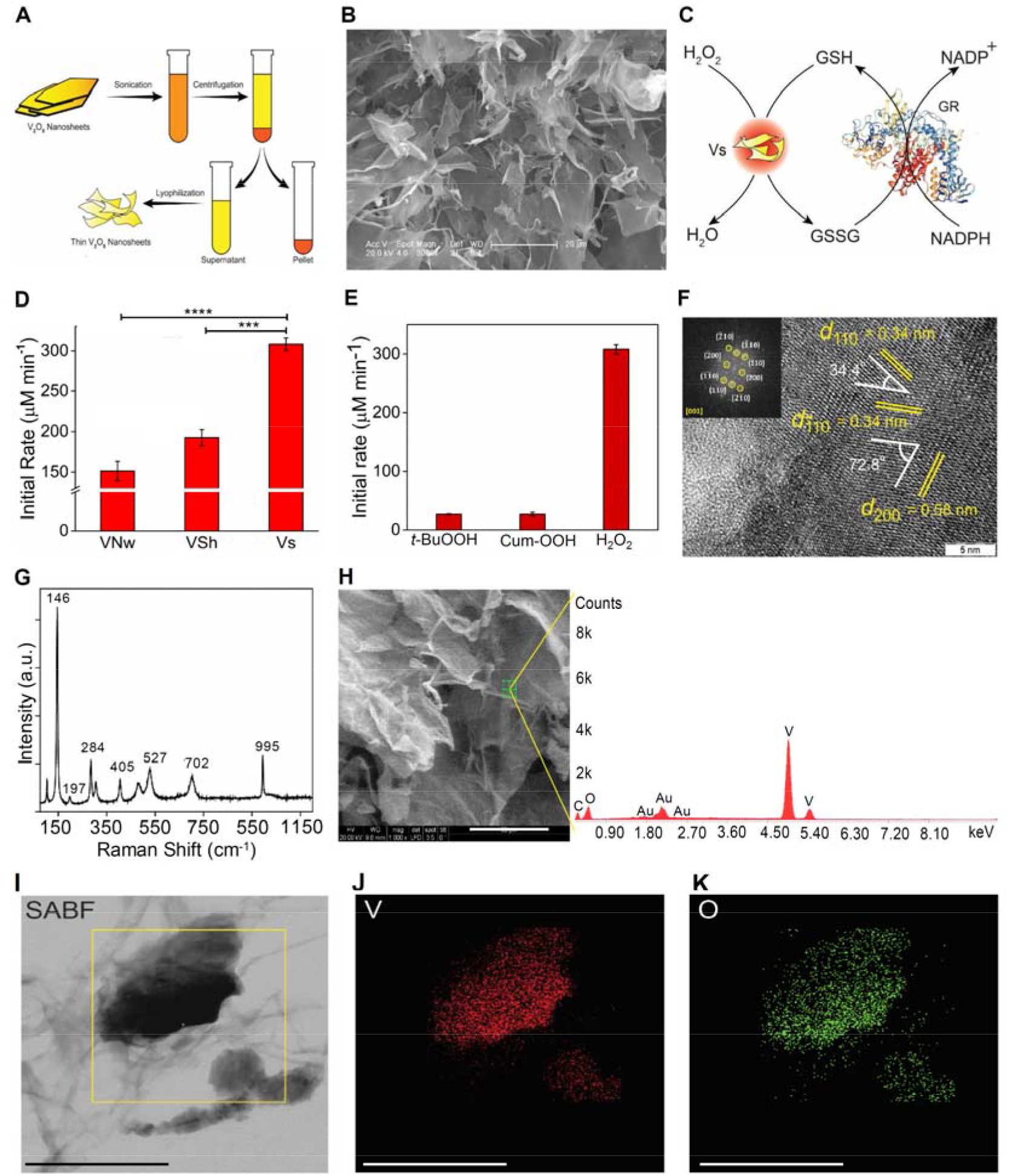
Synthesis and characterization of vanadia (V_2_O_5_) nanoparticles. **(A)** Schematic representation of the synthesis of thin V_2_O_5_ nanosheets (Vs) from VSh. **(B)** SEM image of thin-nanosheets (Vs), scale – 20 µm. **(C)** Schematic representing the glutathione reductase (GR)-coupled assay to measure the GPX-like activity of Vs. **(D)** Comparison of initial activity rate among all three forms of V_2_O_5_ nanomaterials having a common exposed facet [001]. *** P<0.001, **** P<0.0001 by Student’s *t* test. Data is representative of three independent experiment (mean ±SD). **(E)** Bar diagram of the initial rate of Vs with 3 different peroxides, *t*-BuOOH – tertiary-butyl hydroperoxide, Cum-OOH – cumene hydroperoxide and H_2_O_2_. **(F)** High-resolution TEM (HRTEM) and fast Fourier transform (FFT) (inset) of Vs showing the lattice fringes and the exposed plane. **(G)** FT-Raman spectroscopy of Vs showing the peaks corresponding to the orthorhombic phase of the material. **(H)** Energy dispersive spectroscopy (EDS) of Vs. The small peak at 2.0 KeV is due to Au spurting while recording the spectra (Scale – 50 µm). The peak of C is coming from atmospheric carbon. X-Ray mapping images of Vs (Scale – 300 nm). **(I)** Left column: Selective area bright field (SABF) image, **(J)** middle column: distribution of vanadium (V) atoms in red, **(K)** right column: distribution of oxygen (O_2_) atoms in green.

First, we recorded the crystalline nature of the lyophilized, thin nanosheets, Vs, by the powder X-ray diffraction pattern (PXRD) (Fig. S2A). The PXRD pattern was indexed to the standard V_2_O_5_ orthorhombic phase (a = 11.5160 Å, b = 3.5656 Å, c = 4.3727 Å, JCPDS = 41-1426, Space group Pmmn). Second, we examined the crystal facets exposed in the Vs material using high-resolution transmission electron microscopy (HRTEM) and selected area electron diffraction (SAED) pattern analysis. The observed lattice fringes for Vs are (200) and (110) with d-spacing of 0.58 nm and 0.34 nm, respectively, with the interfacial angle of 72.8^°^. The interfacial angle between two planes (110) and (-110) with equidistant d-spacing is 34.4^°^ (Fig. S2B and 1F). These three planes in the HRTEM fall in the common zone axis [001]. In both the figures, SAED patterns (inset) were well indexed along [001] zone axis, which confirms that the surface exposed facets are indeed [001]. These results agree with the most intense peak due to (001) plane observed in the PXRD pattern of Vs.

Third, we performed FT-Raman spectroscopy to determine the nature of bonding formed between the metal and oxygen atoms in the orthorhombic V_2_O_5_ crystals. The FT-Raman spectra showed a peak around 995 cm^-1^, which corresponds to the terminal (V=O) resulting from the unshared oxygen atom of the V_2_O_5_ crystal (Fig. 1G). The peaks detected at lower vibration frequencies are consistent with the lattice vibrations of layered material (*32–34*). We examined the purity of Vs by confirming the detection signal for vanadium (V) and oxygen (O_2_) using point energy dispersive X-ray spectroscopy (EDS) (Fig. 1H). A small signal detected at 2 KeV is due to the gold (Au) sputtering of the sample during spectrum recording. Fourth, we confirmed the elemental composition and purity of the Vs material by selected area bright field (SABF) images and X-ray mapping images (Fig. 1I). Both these techniques confirmed that Vs material has a homogenous distribution of vanadium (V) and oxygen (O_2_) (Fig. 1, J and K). Finally, we determined the oxidation state of the Vs material using X-ray photoelectron spectroscopy (XPS). The analysis revealed binding energies (BE) and full width at half maxima (FWHM) for the V2p3/2 and V2p1/2 peaks as well as the difference in the BE between O1s and V2p3/2 orbitals (12.8 eV). All of this confirms that vanadium exists in the +5-oxidation state in Vs (Fig. S3A – S3C).

### Vs displays efficient H_2_O_2_ linked GPX activity

Having established the biophysical characteristics of Vs, we examined the GPX-mimetic activity of Vs using GR-coupled assay as described earlier (Fig. 1C). Various control experiments, such as reactions lacking GSH/GR/H_2_O_2_, were performed to rule out the possibility of nonspecific reactions. We observed that in the absence of any one of the required constituents, Vs does not show GPX-like activity i.e., no reduction of H_2_O_2_ takes place (Fig. 2A, and S4A). Varying the concentration of Vs from 0 to 20 ng/µL led to a proportional dependence of the initial rate for the reduction of H_2_O_2_ with first-order reaction kinetics (Fig. 2B). Since both H_2_O_2_ and GSH are important for the GPX activity, we performed the activity assay by varying concentrations of H_2_O_2_ (0-400 µM) and GSH (0-7 mM) under steady state condition. Typical enzymatic Michaelis-Menten kinetics was observed for both H_2_O_2_ and GSH (Fig. 2, C and D). The corresponding Lineweaver-Burk plots are depicted in fig. 2E and 2F. For comparison, we simultaneously performed kinetics of Vs, VSh, and VNw. Interestingly, the V_max_ values for Vs and VSh correlate with their surface area with an exception of VNw (Fig. 2, G and H). This is consistent with the differences in the {001} exposed facets among three morphologies. For H_2_O_2_, the K_M_ values obtained for VNw, VSh, and Vs were 44.4 ± 1.7, 57.3 ±3.8, and 112.2 ± 3.8 μM respectively (Fig. 2H). This indicates that the surface of the nanowires and nanosheets (VNw & VSh) are saturated at lower concentrations of H_2_O_2_ (*26*), whereas relatively higher concentrations of H_2_O_2_ are required for the saturation of the surface of thin nanosheets (Vs).

**Figure 2.**
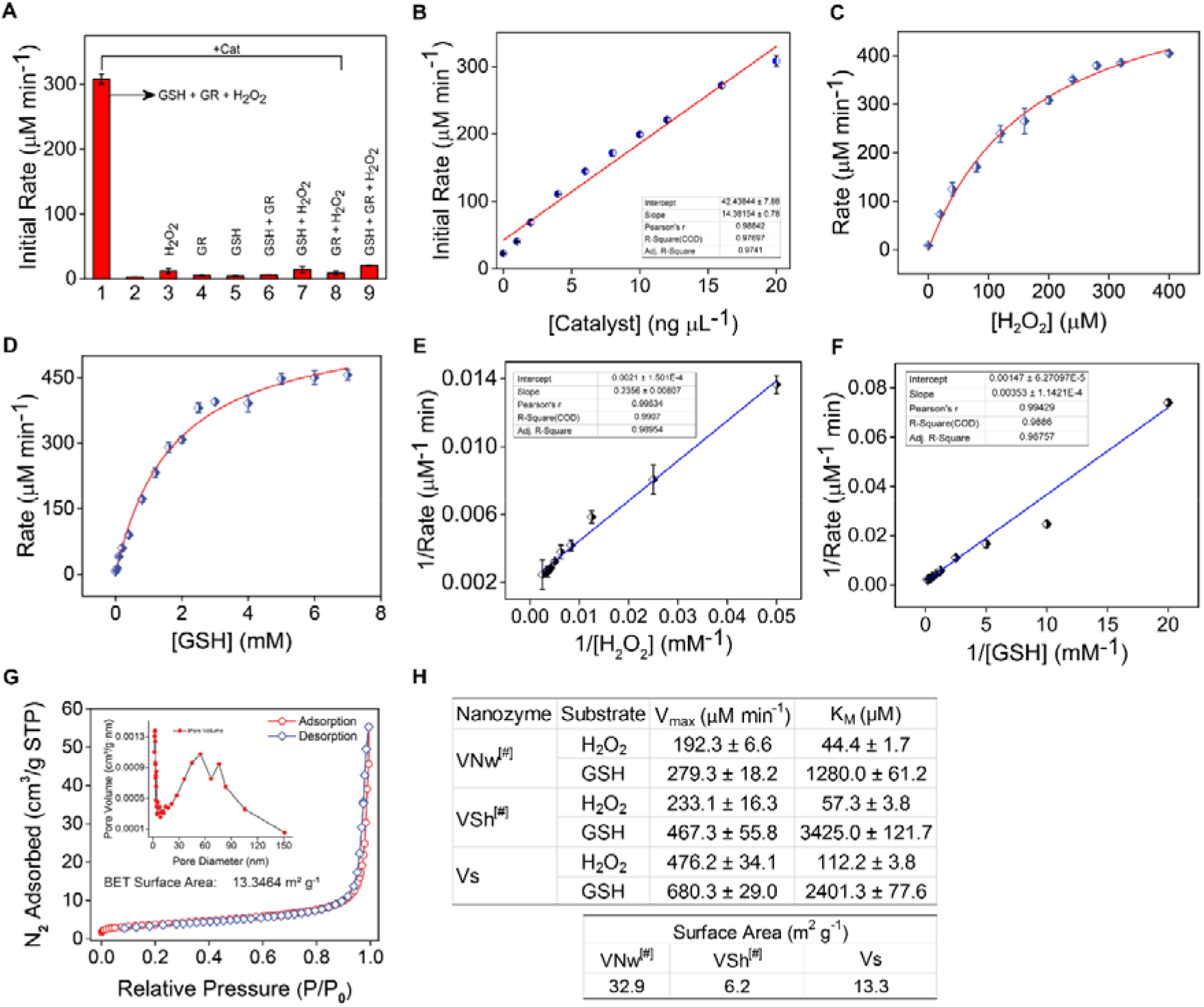
Enzymatic parameters and surface area measurements of Vs. **(A)** Bar diagram shows the initial rate of GPX-like activity of Vs under different assay conditions. **(B)** Dependence of the initial reaction rate for H_2_O_2_ reduction on varying concentration of the catalyst, Vs. **(C)** Michaelis–Menten plot with the variation of H_2_O_2_ (0–400 µM) in the presence of Vs (20 ng/µL), GSH (2 mM), NADPH (0.2 mM), GR (1.7 units) in phosphate buffer (100 mM, pH 7.4) at 25°C. **(D)** Michaelis–Menten plot with the variation of GSH (0–7 mM) in the presence of Vs (20 ng/μL), H_2_O_2_ (200 µM), NADPH (0.2 mM), GR (1.7 units) in phosphate buffer (100 mM, pH 7.4) at 25°C. **(E)** and **(F)** Lineweaver-Burk plot with varying concentration of H_2_O_2_ and GSH in presence of Vs nanozyme respectively. The concentration of NADPH was constant (0.2 mM) in all the assay conditions. **(G)** Surface area measurement by N_2_ adsorption, desorption isotherm and distribution of pore size (Inset). The measured surface area of Vs was 13.3 m^2^/g. **(H)** Enzyme kinetic parameters and BET surface area values of different forms of V_2_O_5_ nanomaterials ^#^ reported from our previous literature (*26*).

The stability of the nanomaterials for the reduction of H_2_O_2_ was examined by performing multiple assay cycles, which demonstrate only a marginal loss of catalytic activity (Fig. S5A). TEM measurements of nanomaterial surface before and after multiple rounds of catalysis indicate no alterations (Fig. S5B and S5C), confirming that Vs performs H_2_O_2_ reduction with unprecedented stability, specificity, and sensitivity *in vitro*.

### Vs mimics GPX activity inside the HIV-1 infected cells

To test the Vs-related GPX activity inside mammalian cells and to understand its influence on HIV-1, we selected the U1 cell line model of HIV-1 latency and reactivation. The U1 cell line is derived from the parent promonocytic cell line U937, wherein two copies of the HIV-1 genome is latently integrated (*35*). The viral replication can be induced by treatment of U1 cells with various pro-inflammatory agents such as phorbol myristate acetate (PMA), tumor necrosis factor-alpha (TNF-α), and granulocyte-macrophage colony-stimulating factor GM-CSF (*35, 36*). We first examined the uptake of Vs nanomaterial by U1 through inductively coupled plasma mass spectrometry (ICP-MS). As shown in fig. 3A, 15 min of Vs exposure led to its buildup inside U1, which was gradually decreased over time such that only a fraction of Vs was retained intracellularly (Fig. 3A). We also found that survival of U1 was not adversely affected by a range of Vs concentration (Fig. S6A).

**Figure 3.**
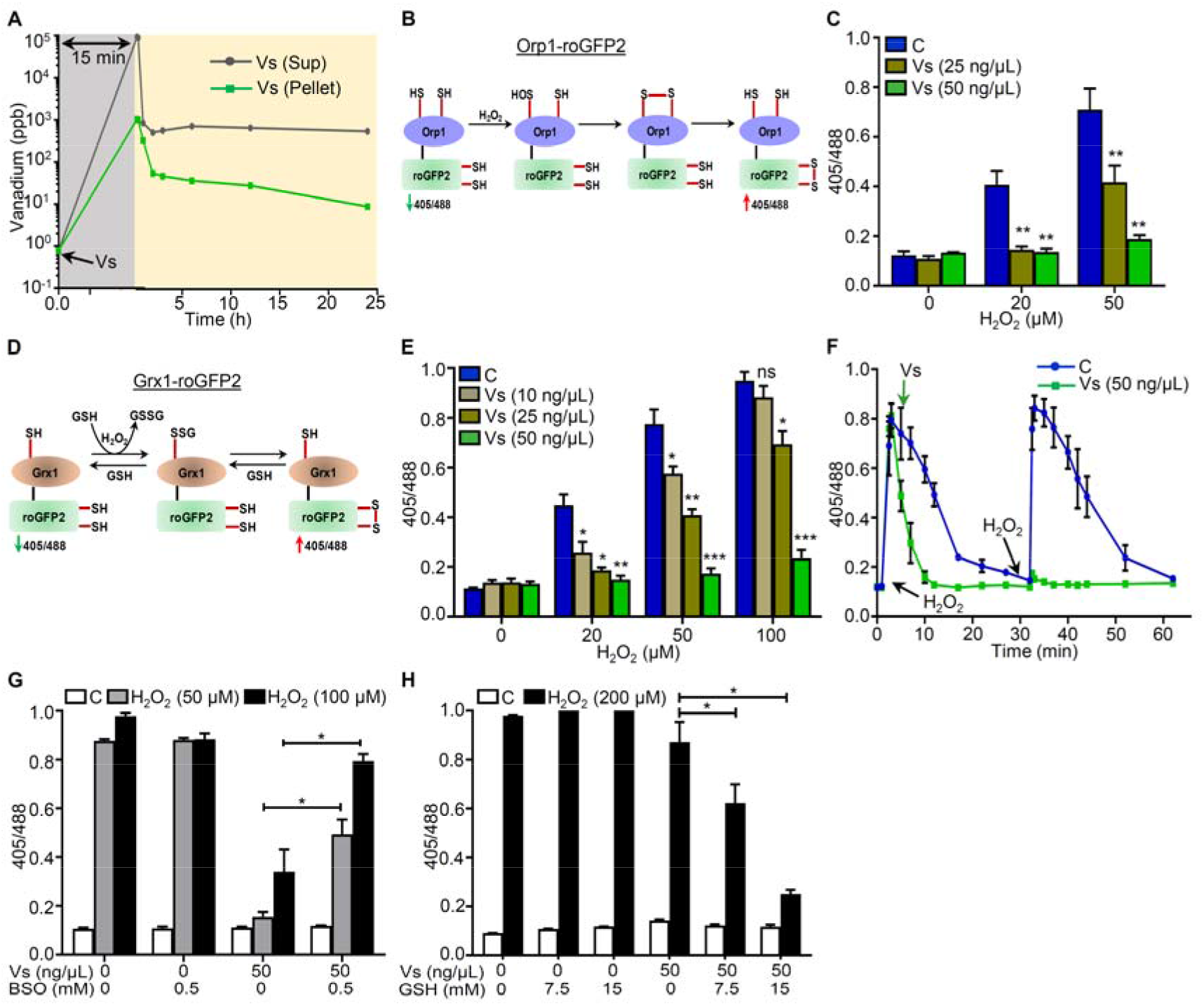
Vs acts as a mimic of GPX in U1 cells. **(A)** U1 cells were treated with 50 ng/ μL of Vs for 15 min (grey area), washed, and kinetics of Vs retention (yellow area) was quantified by measuring cellular and extra-cellular Vs content using ICP-MS. **(B)** Schematic representation showing working principle of Orp1-roGFP2 biosensor. **(C)** U1-Orp1-roGFP2 cells were treated with Vs for 15 min, followed by exposure to H_2_O_2_ for 2 min, and ratiometric response was measured. **(D)** Schematic representation showing working principle of Grx1-roGFP2 biosensor. **(E)** U1-Grx1-roGFP2 cells were pre-treated with Vs for 15 min, exposed to H_2_O_2_ for 2 min, and ratiometric response was measured. **(F)** U1-Grx1-roGFP2 cells were treated with two doses of 50 μM H_2_O_2_ (indicated by the arrows) and the ratiometric response was measured (blue line). Parallelly, U1 cells treated similarly with H_2_O_2_ were exposed to Vs at the indicated time point and the ratiometric response was measured (green line). ****P<0.0001, by Wilcoxon matched-pairs signed rank test. **(G)** and **(H)** U1 Grx1-roGFP2 cells were supplemented with BSO or GSH for 16 h to deplete or replenish GSH, respectively. Following this, cells were treated with Vs for 15 min, exposed to H_2_O_2_, and the ratiometric response was measured. **** P<0.0001, *** P<0.001, ** P<0.01, * P<0.05, by Mann Whitney Test. Asterisks (*) compare Vs treated cells with control cells (C). Data are representative of results from three independent experiments performed in triplicate (mean ± SEM).

To examine the role of Vs in the intracellular reduction of H_2_O_2_ through GPX activity, we exploited Orp1-roGFP2 and Grx1-roGFP2 biosensors that allow non-invasive imaging of the intracellular H_2_O_2_ and *E_GSH_*, respectively (*37–39*). The roGFP2 moiety has two surface exposed cysteines, which undergo disulfide bond formation upon oxidation resulting in an increase in fluorescence excitation intensity at 405 nm along with a relative decrease at 488 nm excitation at a fixed emission of 510 nm (*37*). An inverse relation in 405/488-biosensor ratio was detected upon the reduction of disulfides. The specific equilibration of the roGFP2 dithiol-disulfide redox pair (roGFP2_red_/roGFP2_oxi_) either with H_2_O_2_ or with glutathione redox pair (GSH/GSSG) is efficiently catalyzed by the covalently fused peroxidase Orp1 or glutaredoxin Grx1, respectively (*38, 39*). The fusion of Orp1 with roGFP2 creates a redox relay in which Orp1 mediates near-stoichiometric oxidation of roGFP2 by H_2_O_2_ (*38*) (Fig. 3B). Likewise, Grx1 mediates oxidation of roGFP2 in response to a nanomolar increase in intracellular GSSG upon H_2_O_2_ stress (*37*). The redox relay created by Orp1-roGFP2 or Grx1-roGFP2 demonstrates dynamic behavior as the biosensor ratio returns to basal during recovery from oxidative stress due to normalization of H_2_O_2_ and GSSG levels (*37, 39*).

We created stable transfected U1 cells that express either Orp1-roGFP2 (U1-Orp1-roGFP2) or Grx1-roGFP2 (U1-Grx1-roGFP2) in the cytosol (Fig. S7, A and B). Exposure of U1-Orp1-roGFP2 to H_2_O_2_ for 2 min showed a concentration-dependent increase in the biosensor ratio, consistent with the Orp1-mediated oxidation of roGFP2 by H_2_O_2_ (Fig. 3C). In contrast, pretreatment of U1-Orp1-roGFP2 with 25 and 50 ng/μL of Vs for 15 min diminished biosensor oxidation by H_2_O_2_, consistent with the Vs-catalyzed reduction of H_2_O_2_ (Fig. 3C). Because H_2_O_2_ exposure also leads to oxidation of reduced GSH to GSSG (*9*), we monitored this transformation using Grx1-roGFP2 biosensor (Fig. 3D). The U1-Grx1-roGFP2 cells were challenged with various concentrations of H_2_O_2_ for 2 min, and the sensor response was quantified. We found that the biosensor responds to increasing concentrations of H_2_O_2_ and M of H_2_O_2_ for 2 min results in 90% oxidation of Grx1-roGFP2 (Fig. 3E). The corresponding *E_GSH_* was -240 mV, which is higher than the basal *E*_GSH_ for U1 cells, -320 mV. Pretreatment of U1-Grx1-roGFP2 with Vs for 15 min effectively reduced H_2_O_2_ mediated oxidation of biosensor in a concentration dependent manner (Fig. 3E). Next, we measured the time kinetics of Grx1-roGFP2 oxidation to a low concentration of H_2_O_2_ (50 μM was observed within 2 min of H_2_O_2_ exposure followed by a gradual decrease to the baseline levels in 30 min, indicating efficient mobilization of cellular antioxidant machinery (*37*) (Fig. 3F). In contrast, the addition of 50 ng/μL treatment decreased the biosensor oxidation to baseline levels within 10 min (Fig. 3F). Importantly, a single dose of Vs completely prevented subsequent oxidation of biosensor by H_2_O_2_ (Fig. 3F). This data is fully consistent with earlier results demonstrating multiple cycles of H_2_O_2_ reduction by a single dose of Vs *in vitro*.

Since GPX function is dependent on GSH as an electron donor (*40*), we tested the requirement of GSH in Vs-mediated H_2_O_2_ reduction. We treated U1-Grx1-roGFP2 cells with 0.5 mM buthionine sulfoximine (BSO), which lowers cellular GSH content by inhibiting y-glutamyl cysteine synthetase (GCS) activity (*41*). Following this, cells were treated with Vs for 15 min and exposed to 50 and 100 μM 2 min. As shown earlier, both the concentrations of H_2_O_2_ achieved nearly complete oxidation of Grx1-roGFP2, which was effectively blocked by Vs pretreatment (Fig. 3G). In contrast, pretreatment with BSO attenuated Vs ability to prevent biosensor oxidation by H_2_O_2_ (Fig. 3G). Supplementation of exogenous GSH (15 mM) restored Vs activity as shown by a significant decrease in the biosensor oxidation upon challenge with a saturating concentration of H_2_O_2_ (Fig. 3H).

### Vs subverts HIV-1 reactivation

Studies have shown that H_2_O_2_ treatment reactivates HIV-1 from latency (*15, 42*). Increased oxidative stress was shown to activate the HIV-1 LTR through redox-sensitive transcription factors, such as NF-κB (*11*). On this basis, we reasoned that Vs displaying efficient antioxidant activity could affect redox-dependent reactivation of HIV-1. We first induced HIV-1 expression using low concentrations of PMA (5 ng/mL) and prostatin (1.25 μM), two well-established activators of HIV-1 (*43, 44*). The expression of the HIV-1 *gag* transcript was monitored as a marker of HIV-1 activation by qRT-PCR at various time points post-treatment with PMA/prostatin. Both activators induced HIV-1 transcription with a significant increase observed at 24 h post-treatment (Fig. 4A). Pre-exposure of U1 with Vs or N-acetyl cysteine (NAC-a well-established antioxidant) effectively blocked PMA/prostatin-mediated viral reactivation (Fig. 4, A and B). Using U1-Orp1-roGFP2, we confirmed an increase in the intracellular levels of H_2_O_2_ at 6 and 12 h post-PMA treatment, which was significantly reduced upon Vs pretreatment (Fig. 4C). This indicates that oxidative stress precedes PMA-stimulated virus reactivation and GPX activity associated with Vs counteracted redox-dependent HIV-1 reactivation. The capacity of Vs in lessening HIV-1 activation was also confirmed in a lymphocytic model of HIV-1 latency (J1.1) (Fig. 4D), corroborating that the effect of Vs is not restricted to a cell-type.

**Figure 4.**
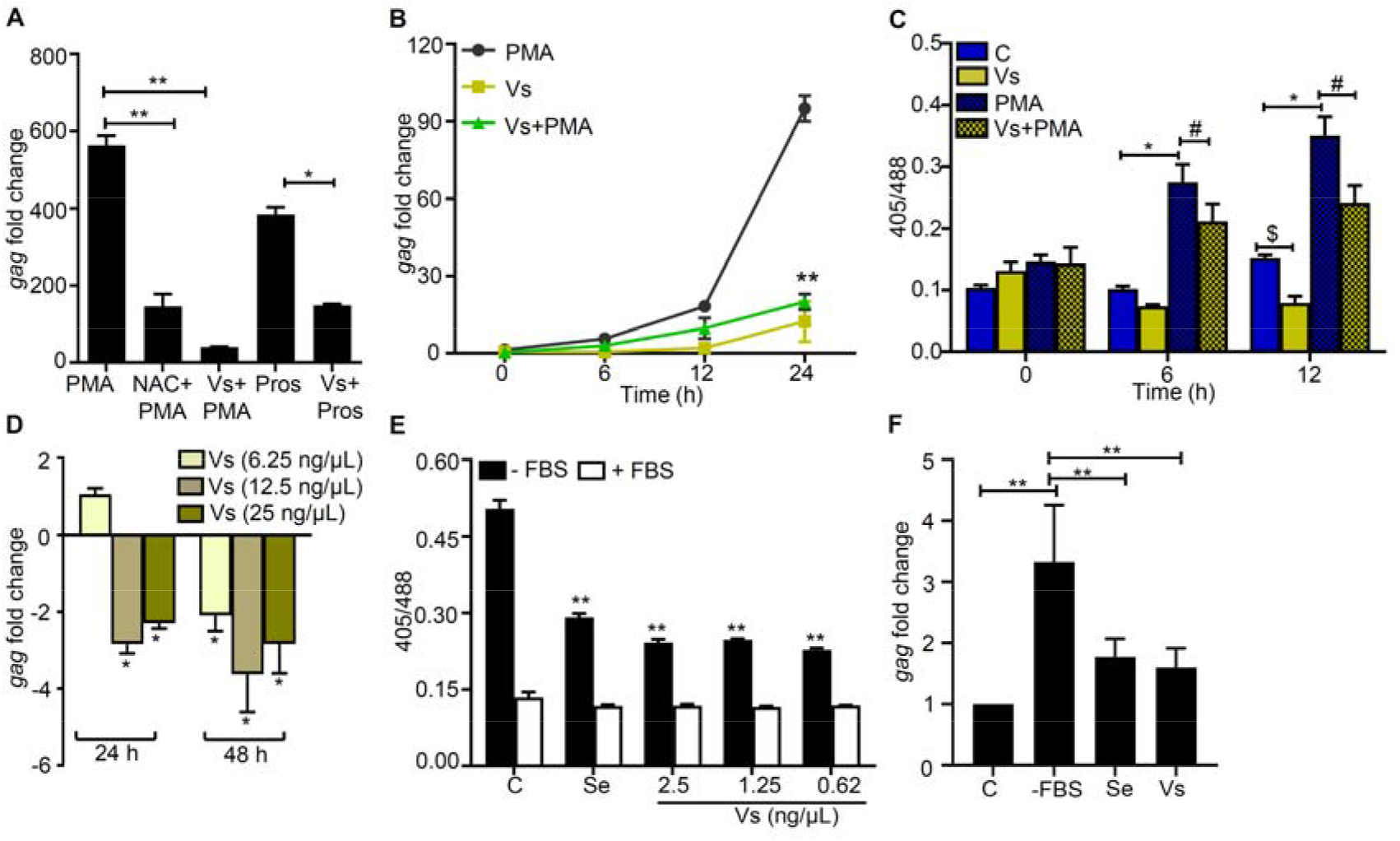
Vs subverts HIV reactivation. **(A)** Vs-treated U1 cells were challenged with 5 ng/mL PMA or 1.25 μM prostratin (Pros) for 24 h and HIV-1 induction was monitored by *gag* RT-PCR. 10 mM NAC, an antioxidant known to subvert PMA mediated viral reactivation, was used as a positive control. **(B)** Vs-treated U1 cells were exposed to PMA and viral activation was measured as a function of time, by *gag* RT-PCR. U1 cells were also treated with Vs or PMA alone. **(C)** Untreated or Vs-treated U1-Orp1-roGFP2 cells were exposed to PMA and the biosensor response was measured at the indicated time points. The biosensor response was also measured for untreated or PMA treated cells. **(D)** J1.1 cells were treated twice with Vs for 15 min at 0 and 24 h time point. HIV-1 induction was measured by *gag* RT-PCR at 24 h and 48 h post-treatment. An untreated control was used for normalization. **(E)** U1-Grx1-roGFP2 cells were serum starved for 30 min in the presence or absence of Vs and sodium selenite (0.5 nM), and the biosensor response was measured. Data were compared to serum starved control cells (C). **(F)** U1 cells were either serum-starved or supplemented with Se (0.5 nM) or Vs (0.62 ng/ μL) and HIV reactivation was measured at 6 h post starvation by *gag* RT-PCR. **P<0.01, $/#/*P<0.05, by Mann Whitney Test. Data are representative of results from three independent experiments performed in triplicate (mean ± SD).

HIV infected individuals suffer from selenium (Se) deficiency that adversely affects the activity of Se-dependent GPX enzyme leading to oxidative stress, HIV reactivation, and exacerbation of disease pathology (*16, 45*). Therefore, Se limitation is a physiologically relevant stimulus that induces oxidative stress and HIV-1 reactivation (*46*). We envisage that Se-independent GPX activity of Vs could replenish the impaired activity of cellular GPX under Se deficient conditions to subvert HIV-1 reactivation. To examine this, we starved U1-Grx1-roGFP2 of fetal bovine serum (FBS; the source of Se) and monitored the change in its antioxidant response over time. We observed an increase in biosensor ratio within 30 min of FBS removal, indicating oxidative stress (Fig. S8A). Supplementation of Vs or Se in the culture medium of Se-deficient U1-Grx1-roGFP2 decreased biosensor ratio, signifying alleviation of oxidative stress by Vs (Fig. 4E). As expected, Se-deficiency triggered HIV-1 reactivation in U1, and addition of Vs or Se had an opposite effect (Fig. 4F).

### Vs adversely affects intracellular replication of HIV-1 and *Mtb*

Along with reactivation, oxidative stress has been shown to promote HIV-1 replication (*9*). Therefore, we next examined the influence of Vs activity on HIV-1 replication. First, we used a stable CD4^+^ T cell clone expressing EGFP (CEM-GFP) under HIV-1 LTR. Infection with HIV-1 significantly induces the expression of CEM-GFP (*47*). The infection of CEM-GFP cells with T cell-tropic HIV-1 provirus (pNL4.3) progressively increased GFP fluorescence over 5 days (Fig. 5A). Addition of 50 ng/μL of Vs for 15 min every 24 h completely blocked GFP expression in the infected CEM-GFP cells (Fig. 5A). We also infected Jurkat CD4^+^ T cells with pNL4.3 virus and measured *gag* transcript, and p24 HIV capsid protein in the whole cell lysate and in the supernatant. Each technique showed a time-dependent increase in HIV-1 replication, while treatment with Vs resulted in a significant inhibition (Fig. 5, B to D). Finally, we infected U937 promonocytic cells with macrophage (M)-tropic HIV pro-virus, pNLAD8, and HIV-1 replication was measured by measuring *gag* transcript at 24 h post-treatment. As shown in fig. 5E, HIV-1 infected U937 showed a 15-fold increase in *gag* transcript, which was reduced to 3-fold in case of Vs pre-treatment. We also examined if antioxidant potential of Vs confers anti-viral response in primary human CD4 T lymphocytes. We pretreated primary CD4 T cells isolated from human peripheral blood mononuclear cells (PBMCs) with Vs (25 ng/µl), infected with HIV-1 NL4.3, and measured HIV-1 replication at 3- and 5-days post-infection. The p24 ELISA confirmed 100 to 200-fold increase in virus replication at 3 to 5 days post-infection, respectively (Fig. 5F). In contrast, pretreatment of Vs restricted virus load to 20- to 80-fold at these time-points (Fig. 5F).

**Figure 5.**
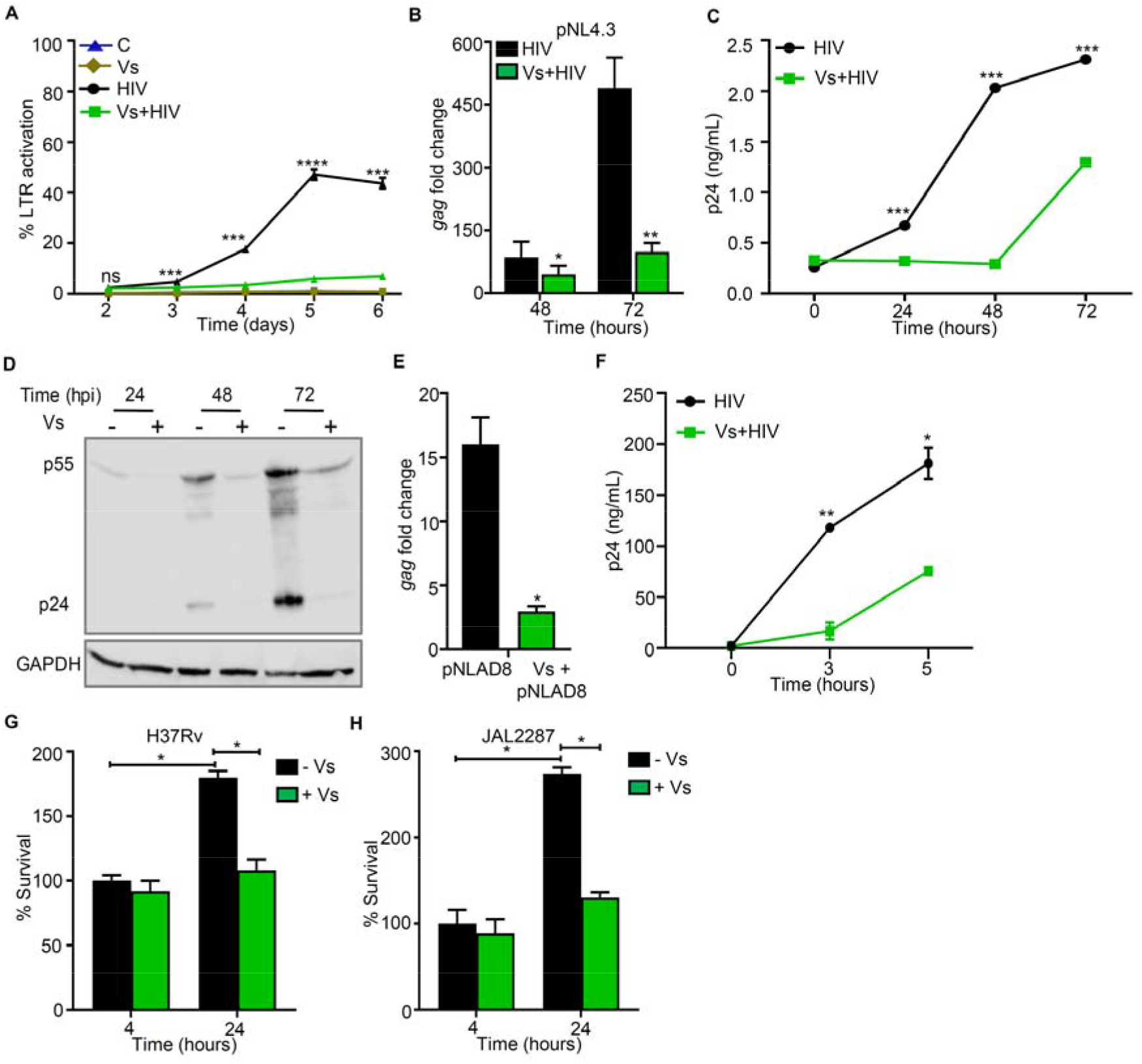
Vs reduces replication of HIV-1 and *Mtb*. **(A)** CEM-GFP cells were pre-treated with 50 ng/ μL of Vs for 15 min and infected with 0.1 moi of T-tropic HIV-1 provirus (pNL4.3) and GFP fluorescence was measured by at 488 nm as an indicator of HIV LTR activity. Vs treatment was repeated every 24 h for the experiment. A similar assay was performed using Jurkat (CD4^+^ T cell line) and viral replication was assessed by **(B)** *gag* RT PCR, **(C)** p24 ELISA in the culture supernatant, and **(D)** immunoblotting for p24 (viral capsid protein) in the whole cell lysate. **(E)** U937 (promonocytes) were pre-treated with 50 ng/μL of Vs for 15 min followed by infection with 1 moi of M-tropic HIV pro-virus (pNLAD8) and viral activation was measured by *gag* qRT-PCR at 24 h post-infection (hpi). **(F)** Primary CD4^+^ T cells purified from human PBMCs were activated, pre-treated with 25 ng/μL Vs for 15 min, and infected with 0.1 moi of T-tropic HIV-1 provirus (pNL4.3). Virus released in supernatant was quantified by p24 ELISA. Vs treatment was repeated every 48 h. Data analysed by Student’s *t* test (mean ± SD). **(G and H)** U1 macrophages pre-treated with 50 ng/μL Vs were infected with drug-sensitive *Mtb* H37Rv or multidrug-resistant JAL2287 strains and bacterial survival was evaluated by enumerating colony forming units (CFUs). Percent survival was calculated by normalizing with CFUs derived from *Mtb* infected Vs-untreated U1 at 4 hpi. All figures except **(A)** and **(C)** were analysed by Mann Whitney test. **(A)** and **(C)** were analysed by 2-way ANOVA with Tukeys multiple comparison test. **** P<0.0001, *** P<0.001, * P<0.05. Data are representative of results from three independent experiments performed in triplicate (mean ± SD).

We have previously shown that oxidative stress promotes synergy between *Mtb* and HIV (*15, 48*). Antioxidant GSH has been shown to reduce *Mtb* load in HIV-infected individuals, while NAC lowers the survival of *Mtb* in mice and guinea pigs (*49, 50*). On this basis, we assessed if the antioxidant potential of Vs affects the survival of *Mtb* in macrophages infected with HIV-1. PMA-differentiated U1 were infected with a laboratory strain of *Mtb* (H37Rv) and a multidrug-resistant patient isolate (JAL 2287) (*51*). Bacterial survival was monitored at 4 h and 24 h post-infection. *Mtb* H37Rv and Jal 2287 displayed 1.78- and 2.74-fold increase in survival at 24 h post-infection, respectively (Fig. 5, G and H). Notably, pretreatment with Vs completely abrogated the ability of *Mtb* strains to multiply inside U1 (Fig. 5, G and H). In sum, Vs catalytic activity efficiently counteracts replication of HIV-1 and *Mtb*.

### Vs dampens the expression of host genes involved in HIV-1 reactivation

Having shown the utility of Vs in counteracting oxidative stress and HIV-1 reactivation, we next examined the underlying mechanisms. We performed expression analysis using the NanoString nCounter system, which permits absolute quantification of multiple RNA transcripts without any requirements for reverse transcription (*52*). We focused on 185 host genes that are known to respond to HIV infection and oxidative stress (Table S1A-S1C). We performed expression analysis on RNA isolated from U1, PMA treated U1, Vs treated U1, and Vs plus PMA treated U1. The fold change (> 1.5-fold, P-value < 0.05) was calculated by normalizing the raw mRNA counts to the geometric mean of the internal control β_2_ microglobulin (B2M).

A total of 123 genes showed differential expression under the conditions tested (Fig. 6A). Overlap analysis confirmed 55 of 123 genes to be common in each category (Fig. 6B). Treatment with PMA induced the expression of genes associated with ROS and RNS (reactive nitrogen species) generation (*e.g.,* NADPH oxidase subunits [NCF1, NCF2] and nitric oxide synthase [NOS2]). Genes involved in antioxidant response, including catalase (CAT) and superoxide dismutase 2 (SOD2), were downregulated upon PMA treatment (Fig. 6C). Overall, these changes are consistent with increased oxidative stress in response to PMA triggered HIV-1 reactivation (*15*). Up-regulation of genes involved in reducing free iron pool (ferritin heavy chain1; FTH1), maintaining GSH balance (cysteine/glutamate transporter; SLC7A11), (Fig. 6C), indicate a compensatory mechanism to protect from oxidative conditions induced by PMA (*53, 54*). The transcription factor Nrf2 is the major activator of antioxidant systems (*55*). Surprisingly, a majority of Nrf2 dependent antioxidant systems such as GSH biosynthesis/recycling (*e.g.,* GSS, GPX1, GPX4, GSTP1), thioredoxins (*e.g.,* TXNRD2), and peroxiredoxins (PRDX6) were down regulated upon treatment with Vs alone or Vs plus PMA (Fig. 6C), indicating an adverse influence of Vs on U1 cells. One likely possibility is that the natural antioxidant defense mechanisms are attenuated by a feedback-like mechanism because of the potent antioxidant properties of Vs. Also, the expression of superoxide producing system (NCF1 and NCF2) was repressed in Vs alone and Vs plus PMA treated U1 (Fig. 6C), which can further reduce intracellular ROS levels.

**Figure 6.**
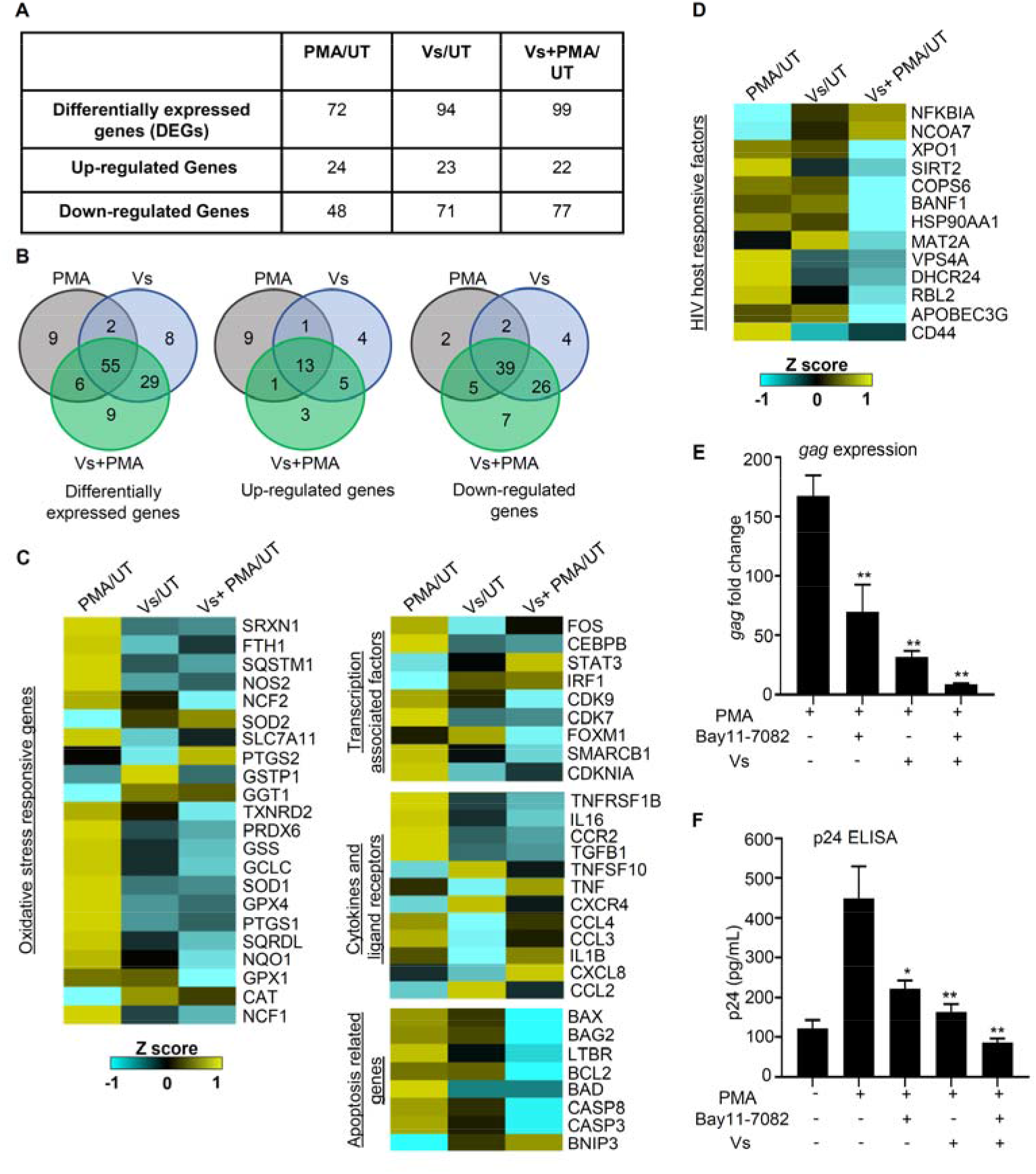
Expression analysis of host pathways associated with HIV-1 activation. U1 cells were treated with Vs (50 ng/ μL) for 15 min, followed by PMA treatment for 12 h. Total RNA was isolated from untreated (UT), PMA-treated, Vs-treated and Vs+PMA treated U1 cells and expression analysis of 185 genes specific to oxidative stress and HIV host response was measured using the NanoString technology. **(A)** Number of differentially expressed genes (DEGs) under the three comparison conditions i. PMA/UT, ii. Vs/UT and iii. Vs+PMA/UT. **(B)** The venn diagram showing overlap of DEGs significantly perturbed under different comparison conditions. **(C and D)** Heat map showing multiple categories of DEGs under PMA/UT, Vs/UT and PMA+Vs/UT comparisons. mRNA counts were normalised using the internal control β_2_ microglobulin (B2M), and fold change (FC) was calculated using the nSolver 4.0 software. Genes showing an absolute FC >1.5, and P <0.05 were considered as significantly altered. **(E and F)** Vs-treated or untreated U1 cells were exposed to Bay11-7082 (7.5 µM). PMA mediated HIV reactivation at 12 h was monitored by *gag* RT PCR and p24 ELISA. **P<0.01, *P<0.05, by Mann Whitney Test. Data are representative of results from three independent experiments performed in triplicate (mean ± SEM). Asterisks (*) compare different treatment conditions with PMA-treated cells.

Genes known to be associated with HIV-1 activation such as transcription factors (*e.g.,* FOS and CEBPB) (*56, 57*), inflammatory cytokines/receptors (TGF β1, TNFRSF1B, and IL16) (*58–60*), and chemokines (CCL3 and CCL4) (*61*) were induced upon PMA treatment and repressed by Vs plus PMA (Fig. 6C). Several genes encoding proteins associated with HIV-1 replication, packaging, budding, and fitness (*e.g.,* APOBEC3G, CD44, XPO1, VPS4A, DHCR24) were down-regulated upon Vs plus PMA treatment as compared to PMA alone (Fig. 6D). It is known that cells latently infected with HIV-1 are refractory to apoptosis, whereas increased apoptosis promotes HIV-1 reactivation (*62*). Consistent with this, a majority of genes encoding pro-apoptotic proteins (*e.g.,* BAD, BAX, CASP3, and CASP8) were substantially repressed upon Vs plus PMA treatment as compared to PMA alone (Fig. 6C). In addition, a cellular inhibitor of transcription factor NF-κB (i.e NFKBIA) was highly induced upon Vs or Vs plus PMA treatment (Fig. 6D). Since NF-κB is known to activate HIV-1 in response to oxidative stress (*10*), the induction of its inhibitor (NFKBIA) by Vs is indicative of reduced reactivation of HIV-1 through NF-κB. Based on this, we hypothesize that a well-established pharmacological inhibitor of NF-κB (*E*)3-[(4-methylphenyl)sulfonyl]-2-propenenitrile (BAY11-7082) (*63*) would synergies with Vs to efficiently subvert HIV-1 reactivation. To examine this, we exposed U1 cells pretreated with Vs to BAY11-7082 and HIV-1 reactivation in response to PMA was monitored by measuring the levels of *gag* transcript and p24 capsid protein. Consistent with our hypothesis, exposure of U1 to both Vs and BAY11-7082 suppresses reactivation of HIV-1 which supersedes that produced by either Vs or BAY11-7082 alone (Fig 6E and 6F). Overall, Vs not only affected the expression of redox pathways but as a consequence also modulates the expression of pathways coordinating the inflammatory response, viral fitness, transcription, and apoptosis to subvert HIV-1 reactivation.

### Vs mitigates oxidative stress and HIV-1 associated pathologies *in vivo*

Given our findings that Vs diminishes oxidative stress to reduce HIV-1 reactivation in cell lines, we sought to determine the impact of Vs on oxidative stress and HIV-1 associated pathologies *in vivo*. We used an HIV-1 transgenic mouse (Tg26), which harbors a replication-incompetent, Δ*gag* variant of pNL4.3/HIV (*64*). Expression of HIV-1 proteins in Tg26 promotes oxidative stress and recapitulates clinical lesions observed in HIV infected patients (*e.g.,* lung damage and nephropathy) (*64–66*).

We treated Tg26 C57BL/6 mice with Vs (1 mg/kg body weight) intra-peritoneally (i.p.). Untreated or Vs-treated animals were euthanized 3 h post-treatment and oxidative stress was measured in the lungs (Fig. 7A). The lung homogenates were stained with redox-active fluorescent dye 2’,7’ – dichlorofluorescein diacetate (DCFDA), and subjected to flow cytometry. As shown in figure 7B, total cellular populations within lung homogenates derived from Vs treated Tg26 displayed diminished fluorescence as compared untreated Tg26. Based on the forward and side scatter gating, we gated different populations representing mostly granulocytes (P2), monocytes (P3) and lymphocytes (P4). The level of fluorescence within multiple cell types was consistently lower in the case of Vs treated Tg26 than untreated Tg26 animals (Fig. 7B). Using antibodies against CD64 and MERTK markers, we specifically gated lung macrophages and confirmed lower ROS in cells derived from Vs treated Tg26 animals than untreated Tg26 (Fig. 7C). The lung cells derived from *Wt* C57BL/6 showed ROS levels comparable to Vs treated Tg26 animals (Fig. 7, B and C). Consistent with a strong antioxidant activity of Vs, expression of other cellular antioxidant genes were either down-regulated (*e.g,* GPX1) or unchanged (GPX4, SOD1, and SOD2) in the lungs of Vs treated Tg26 as compared to untreated animals (Fig. 7D and Fig. S9A). One of the clinical signs of HIV infection is inflammation (*67*). The Tg26 mice display edema in various organs, which is a marker for inflammation and fluid accumulation (*65*). Treatment with Vs significantly reduced lung edema in Tg26 animals (Fig. 7E). Lastly, we confirmed that Vs treatment reduced the expression of HIV-1 transcript (*tat-rev* region) by ∼65% in Tg26 mice (Fig. 7F). Altogether, our findings position Vs as a potential alternative to natural GPX enzyme(s) for regulating redox signaling, inflammation, and virus expression *in vivo*.

**Figure 7.**
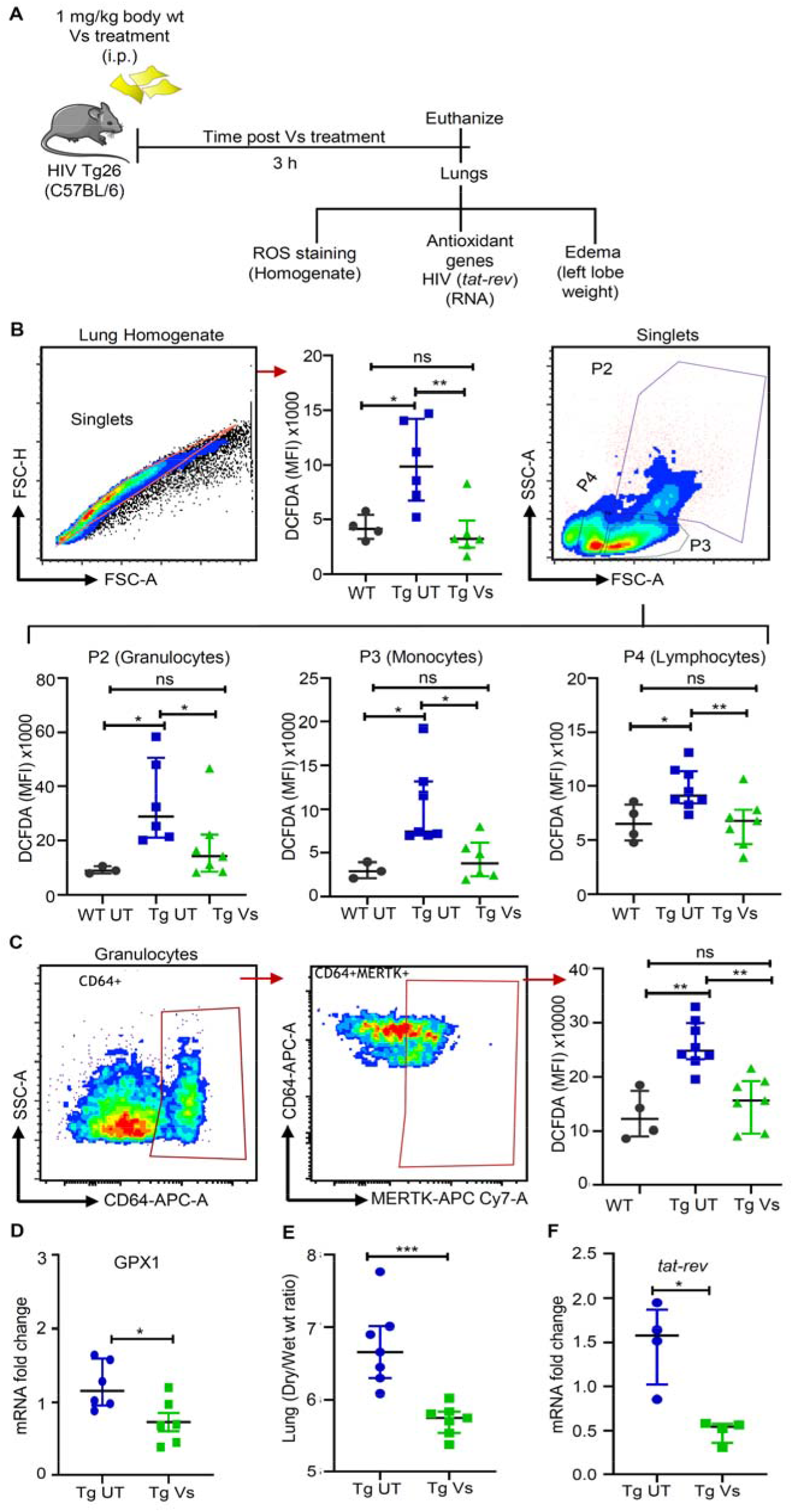
Vs reduces oxidative stress, edema, and HIV expression *in vivo*. **(A)** Strategy to investigate the antioxidant function of Vs in HIV Transgenic mice, Tg26. HIV Tg26 (n=8), were left untreated or treated with 1 mg/kg Vs for 3 h. Untreated WT C57BL/6 was used as a control. **(B)** Lung homogenates were stained with DCFDA and analysed by flow cytometry for ROS detection. Singlets, followed by different cell clusters comprising mainly of granulocytes (P2), monocytes (P3) and lymphocytes (P4) were gated from the whole lung homogenate and ROS levels were plotted as median fluorescence intensity (MFI). **(C)** Pan-macrophage surface markers, CD64 and MERTK, were used to stain total lung macrophage population, ROS levels were quantified by DCFDA staining and plotted as MFI.**(D)** RNA was isolated from the right lung lobe and expression of GPX1 in Tg UT and Tg Vs animals was checked by qRT-PCR. GPX1 expression values were normalised with Tg UT. **(E)** Lung edema plotted as the ratio of wet versus the dry weight of left lobe. **(F)** Expression of *tat-rev* region as a marker for HIV-1 transcription was evaluated by qRT-PCR using RNA from lungs of Tg UT and Tg Vs animals. Data has been plotted as median ± interquartile range. *** P<0.001, ** P<0.01, * P<0.05, by Mann Whitney Test.

## Discussion

Studies exploring the application of antioxidant nanozymes in targeting human pathogens are limited. In-depth cellular studies using laboratory models that mimic the physiological environment during infection could help predict the clinical potential of nanozymes and will encourage new designs that are efficacious in humans. Using several biophysical, biochemical, cellular, genetic biosensors, animal model, and expression technologies, we demonstrated the biomedical application of V_2_O_5_ based nanozymes in counteracting redox stress and reactivation of HIV-1. We showcase the remarkable efficacy of Vs against various physiological conditions, including Se starvation and infection with secondary pathogens such as *Mtb*.

V_2_O_5_ nanoparticles possess intrinsic peroxidase-mimicking activity (*8*). By modifying V_2_O_5_ nanomaterial into various morphologies, we reported efficient GSH-dependent GPX activity of V_2_O_5_ nanozymes (*26*). In the present study, we further fine-tuned the crystal facets of V_2_O_5_ nanomaterial to generate a thinner nanosheet (Vs), which displayed high catalytic activity in the presence of GSH. Other properties, such as the formation of stable V-peroxo species that ensure GSH specificity, multiple recycling without loss of activity, and stability in a range of organic solvents (698) motivated us to exploit V_2_O_5_ nanomaterial in understanding GSH/GPX based redox signaling during HIV-1 infection. Until now, studies examining the antioxidant function of nanozymes relied on *in vitro* enzymatic assays or chemical analyses of redox metabolites (*e.g.,* GSH/GSSG) in whole cells or tissues. These invasive methods introduce oxidation artifacts and preclude observation of real-time changes in redox physiology upon nanozyme treatment. We circumvented these issues by applying non-invasive genetically encoded biosensors of H_2_O_2_ (Orp1-roGFP2) and *E_GSH_* (Grx1-roGFP2) to dynamically assess the activity of Vs in reducing intracellular H_2_O_2_ and maintaining GSH homeostasis.

Numerous studies have indicated a link between GPX activity and HIV-1 *in vitro* and *in vivo* (*16, 21*). Remediation of H_2_O_2_ by GPXs potently reversed NF-κB-mediated HIV-1 transcription (*21*). We have demonstrated that latently infected cells efficiently metabolized H_2_O_2_ likely via endogenous GPXs (*15*). Expression of GPXs was elevated in monocytes and lymphocytes harboring latent HIV-1, whereas expression was diminished in cells and in PBMCs of patients during active HIV-1 replication (*15*). Since the activity of cellular GPXs is selenium (Se)-dependent, studies have found a high correlation between Se-deficiency, GPX activation, and HIV-related mortality (*45*). Importantly, HIV-1 infected T-cells showed the general downregulation of cellular Se-proteins (*69*). We found that Se depletion induced the oxidative shift in *E_GSH_* and promotes HIV reactivation. Importantly, Vs supplementation under Se depleted conditions was sufficient to diminish oxidative stress and HIV-1 reactivation. Mechanistically, our Nanostring data confirm that GPX-like activity of Vs efficiently suppresses the expression of redox-dependent transcription factors, pro-inflammatory cytokines/chemokines, and pro-apoptotic molecules required for HIV-1 reactivation. In line with our findings, transcriptomics of whole blood RNA from elite controllers (ECs) to that of individuals with high viral load showed that cytokine signaling, apoptosis, anti-viral response, and immune activation are mainly affected (*70–72*). Our data align well with the meta-analyses where pathways promoting oxidative stress and viral transcription such as NCF2, ACTB, FOS, CEBPB are downregulated in ECs and antioxidants (GPX1 and GPX4) were induced as compared to viremic progressors (*71, 72*).

Individuals co-infected with HIV and *Mtb* pose major problems in disease management. We showed that basic mechanisms such as intracellular redox potential and bioenergetics contribute to synergy between HIV and *Mtb* (*15,48,73*). Diminished levels of host GSH, GPX4, and increased ROS contribute to the survival of *Mtb in vivo* (*74*). Similar to these observations, supplementation of GPX activity by Vs hampered multiplication of *Mtb* and drug-resistant clinical isolate in U1 cells pre-exposed to Vs. Finally, the biological importance of our findings comes from the data showing reduced oxidative stress, inflammation, and expression of HIV-1 transcripts in Tg26 mice treated with Vs. While our findings provide a clear relationship between catalytic activity of Vs with HIV-1 reactivation and redox signaling, future experiments are needed to exploit Vs for the development of sustainable, affordable, and safe therapeutics against HIV-1 infection. The importance of nanomaterials in infectious diseases is emerging as silver and gold-based nanomaterials have been shown as promising candidates for therapeutic applications against bacterial and viral pathogens (*75–78*).

In conclusion, we have presented compelling experimental evidence that strongly suggests the application of Vs in modulating redox signaling associated with HIV-1 latency and reactivation. Antiretroviral therapy (ART), while effective in reducing viral load, remains unsuccessful in eradicating HIV from latent reservoirs and induces systemic oxidative stress (*79*). Recently, a concept called “block and lock” showed that blocking host (*e.g.,* heat shock protein 90 (HSP90), NF-κB, protein kinase C (PKC) or viral factors (*e.g.,* Tat) involved in reactivation significantly delayed viral rebound upon interruption of ART (*80*). Our findings on Vs showed promise in preventing reactivation from latency, raising the possibility of locking HIV in an extended latency by blocking redox-mediated HIV-1 transcription, and also improving the health of individuals on ART by minimizing oxidative stress.

## Materials and methods

### Preparation of thin vanadia (V_2_O_5_) nanosheets (Vs)

Thin V_2_O_5_ nanosheets (Vs) were synthesized from the crude V_2_O_5_ nanosheets (VSh). Briefly, 2 mM of V_2_O_5_ powder was dispersed in 15 ml ultrapure water for 20 min. Then 15 mL H_2_O_2_ (30% w/v) was added dropwise. During the addition of H_2_O_2_ the colour of the solution changed from yellow to orange and then to red. The red solution turns dark brown after stirring for 2.5 h at room temperature (RT). This reaction was strictly performed in a fume hood due to its exothermic nature. After continuous stirring for 2.5 h, 10 mL of ultrapure water was added into the mixture and heated to 60°C overnight to form a brownish gel (V_2_O_5_.nH_2_O). This gel was dried at 100°C for 12 h and subsequently calcined at 400°C for 2 h to get crude VSh. Following this, VSh was probe sonicated in ultrapure water for 2 h to get a dense dispersion of nanosheets. The dispersion was then centrifuged at 3000 rpm for 5 min and the bright yellow colour supernatant was lyophilised to obtain the powdered form of thin V_2_O_5_ nanosheets (Vs).

### Characterization of Vs

Powder X-ray diffraction (PXRD) was recorded by Phillips PANalytical diffractometer using a CuK α(= λ 1.5406 Å) radiation. The emission current and accelerating voltage used in the diffractometer were 30 mA and 40 kV respectively. For morphological and elemental characterization, EDS and scanning electron microscopy (SEM) were performed on FEI Sirion UHR SEM and ESEM-Quanta respectively. Transmission electron microscopy (TEM), High resolution transmission electron microscopy (HRTEM), and X-ray mapping images were recorded on JEOL transmission electron microscope operated at 200 kV after casting a drop of nanoparticle dispersion in isopropyl alcohol, over a Cu grid. FT-Raman spectra were recorded using a Renishaw in-Via Raman Microscope (Renishaw Inc, UK), with excitation wavelength 514 nm. To perform all the enzyme mimetic activity assay, SHIMADAZU UV-2600 spectrophotometer was used. X-ray photoelectron spectroscopy (XPS) was performed using AXIS Ultra, KRATOS ANALYTICAL, SHIMADAZU. The surface area measurement was performed by Brunauer-Emmett-Teller (BET) method on the micromeritics surface area analyzer model ASAP 2020.

### GPX – mimicking activity of V_2_O_5_ nanoparticles (NPs)

The GPX-like activity of V_2_O_5_ NPs was assessed spectrophotometrically by using the standard GR-coupled GPX assay (*8*). The components and the concentration used in this assay mixture were GSH (2.0 mM), NADPH (0.2 mM), GR 1.7 U, catalyst 20 ng/µL, and H_2_O_2_ (0.2 mM) in sodium phosphate buffer pH 7.4 at 25°C. The rate of the reaction was quantified by following the decrease in the absorbance of NADPH (ε = 6220 M cm at 340 nm) to form NADP which is equal to the rate of conversion of H_2_O_2_ to H_2_O.

### Dynamic response of U1-Grx1-roGFP2 cells towards oxidative stress

Oxidation-reduction kinetics of the Grx1-roGFP2 biosensor were measured by flow cytometry, as demonstrated earlier (*15*). Briefly, the basal redox state of 1 × 10^6^ U1-Grx1-roGFP2 cells was measured, following which 50 μM H_2_O_2_ was added after 2 min. Biosensor oxidation and the kinetics of its subsequent recovery were monitored. Parallelly, Vs was added to a set of H_2_O_2_ treated cells at the point of maximum oxidation, and recovery of the biosensor in the presence of Vs was noted. After complete recovery of the cells from oxidative insult, both untreated and Vs treated cells were challenged with another bolus of 50 μM H_2_O_2_, and the biosensor dynamics were monitored by flow cytometry. Percentage oxidation of the Grx1-roGFP2 biosensor was determined by equating maximal oxidation by 10 mM H_2_O_2_ as 100%.

### Determination of the specificity of Vs towards GSH

The specificity of Vs towards GSH as cofactor was determined by modulating cellular GSH levels. U1-Grx1-roGFP2 cells were treated with 0.5 mM of BSO, an inhibitor of GSH biosynthesis, or supplemented with 7.5 or 15 mM GSH for 16 h. Following this, the cells were treated with 50 ng/μL of Vs and challenged with various H_2_O_2_ concentrations for 2 min. The biosensor response in U1 cells was measured by flow cytometry.

### Assessing the effect of Vs on active HIV replication

CD4^+^ T cell line – CEM-GFP and Jurkat- and monocytic cell line U937 were infected with laboratory adapted HIV-1 strains pNL4.3 (T–tropic) and pNLAD8 (M– tropic), respectively. 0.5 × 10^6^ untreated or Vs treated cells of each type were suspended in 500 μL of Opti MEM media and infected at multiplicity of infection (moi) 0.1. The cells were incubated at 37ºC for 4 h and mixed intermittently during the infection period. After 4 h, the cells were washed to remove unbound virus and supplemented with complete RPMI media with 10% FBS. Vs treatment was repeated every 24 h for Jurkat and CEM-GFP cells. CEM-GFP cells were grown till 5 days and LTR activation was assessed from day 2 to day 5 post-infection by flow cytometry. Viral replication in Jurkat and U937 cells was monitored by qRT-PCR at indicated time points.

### Nanostring gene expression analysis

Expression levels of 185 genes responsive to oxidative stress and HIV infection were analysed in untreated U1 cells, U1 cells treated with PMA or Vs alone, and a combination of Vs plus PMA. The Nanostring nCounter analysis system was utilized for this purpose. Briefly, the assay was performed with 100 ng of total RNA, isolated from untreated or treated cells using the Qiagen RNeasy kit. The purity of the RNA was confirmed spectrophotometrically using Nanodrop Lite Spectrophotometer (Thermo Scientific). The nCounter probes are barcoded DNA oligonucleotides complementary to the target mRNA. Hybridization and counting were performed according to the manufacturer’s protocol (*52*) using a customized panel of 185 genes. 6 housekeeping control genes were included in the panel. Data analysis was done using nSolver 4.0. B2M was used as an internal control due to its minimum % CV.

### Selenium starvation and HIV-1 reactivation

U1-Grx1-roGFP2/U1 cells grown in complete RPMI medium were harvested and washed three times with serum-free RPMI to remove traces of Se. Cells were seeded in 24 well plates and incubated for 30 min, 1 h, and 2 h in serum free medium. Parallelly, the cells were treated with various concentrations of Vs for 15 min and cultured as mentioned above. 0.5 nM sodium selenite (Se source) was used as a positive control. Cells were harvested at indicated time points and the biosensor response was measured by flow cytometry. Viral reactivation and expression of host antioxidant genes were analysed 6 h post starvation, as mentioned earlier.

### Methods included in the supplementary information

Dispersion of V_2_O_5_ NPs, treatment of cell lines with Vs, internalization of Vs by U1, mammalian and bacterial Cell Culture, preparation of stable cell lines and validation using flow cytometry, assessment of Vs antioxidant activity and redox potential measurement, propidium iodide (PI) staining, HIV reactivation in U1 cells and qRT-PCR analysis, p24 detection by Immunoblotting and ELISA, bacterial Survival assays in U1 cells, isolation and infection of primary CD4^+^ T cells, animal experiments with HIV-Tg mice, pulmonary edema analysis, ROS staining of lung homogenates.

### Statistical analysis

All statistical analyses were performed using the GraphPad Prism software (Version 8.1). The data values are indicated as mean ± S.D. Statistical significance between two non-parametric test groups was determined using the Mann Whitney Rank Sum test, unless specified. Analysis of Nanostring data was performed using the nSolver platform. Differences in *P* values <0.05 were considered significant.

## Acknowledgements

This work was supported by Wellcome Trust-Department of Biotechnology (DBT) India Alliance grant IA/S/16/2/502700 (A.S.) and in part by DBT grants BT/PR11911/BRB/10/1327/2014, BT/PR13522/COE/34/27/2015 and BT/HRD/NBA/39/07/2018-19 (A.S.), DBT-IISc Partnership Program grant 22-0905-0006-05-987 436, and the Infosys Foundation. A.S. is a senior fellow of Wellcome Trust-DBT India Alliance. G. M. acknowledges the DST Nano Mission (SR/NM/NS-1380/2014) and SERB (SB/S2/JCB-067/2015), DST, New Delhi for funding. SS and SG acknowledge fellowships from the University Grants Commission (UGC) and Indian Institute of Science (IISc) respectively. We thank Dr. Amit A. Vernekar for helpful discussions. We gratefully acknowledge the NanoString services provided by TheraCUES Innovations Pvt Ltd, Bangalore. We also thank the AFMM Facility, CeNSE, IISc for the microscopic and spectroscopic facilities.

## Supplementary Material

### List of Supplementary Materials

Fig. S1. SEM and initial reaction rate for Vanadia Nanowires and crude nanosheets.

Fig. S2. XRD pattern and TEM of Vs

Fig. S3. X-Ray Photoelectron Spectroscopy (XPS) of Vs.

Fig. S4. GR- coupled assay for Vs under varied conditions.

Fig. S5. In vitro recycling ability of Vs.

Fig. S6. Survival assay for Vs-treated U1 cells.

Fig. S7. Preparation of Stable cell lines.

Fig. S8. Time dependent induction of Selenium (Se) starvation-mediated oxidative stress.

Fig. S9. Antioxidant gene expression in lungs of Vs treated Tg26.

Table S1. List of differentially expressed genes from U1 cells either left Untreated (UT), or treated with PMA, Vs or Vs+PMA.

Table S2. List of primers used in the study.

Supplementary Experimental procedures

### Supplementary Figures

**Figure. S1.**
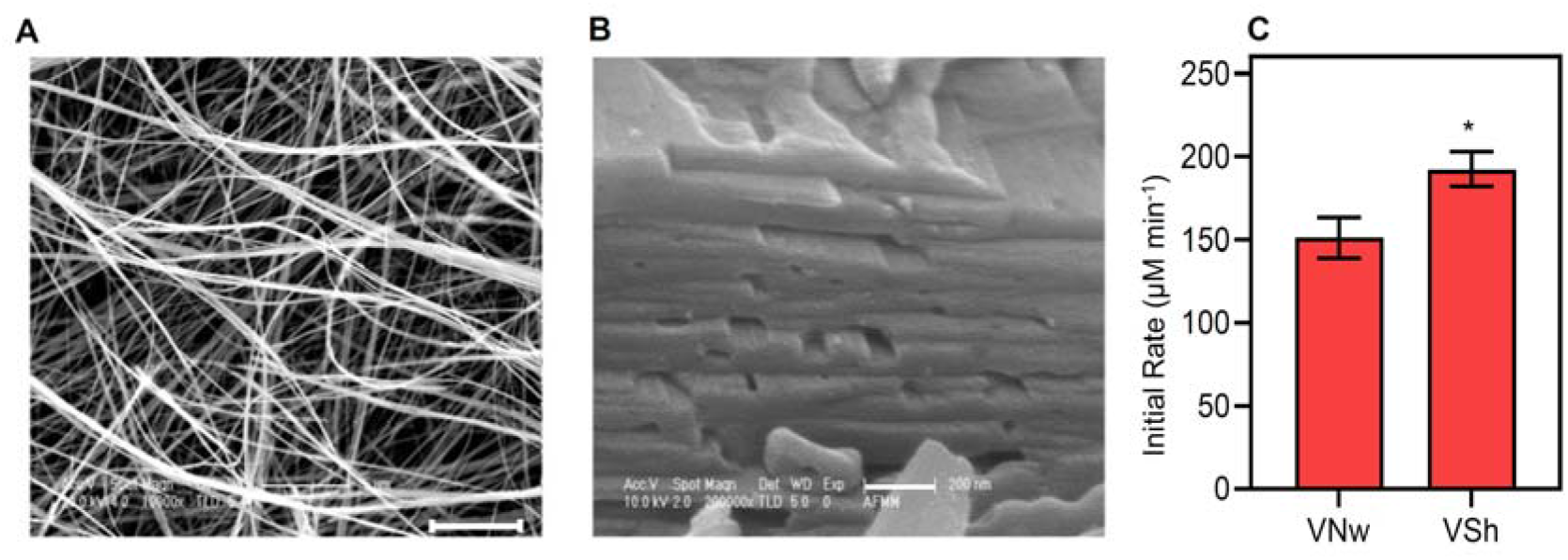
SEM and initial reaction rate for Vanadia Nanowires and crude nanosheets. Scanning electron microscopy (SEM) images of **(A)** nanowires (VNw; scale – 5 µm) and **(B)** crude nanosheets (VSh; scale – 200nm). **(C)** Bar graph shows the initial rate of activity for VNw and VSh. * P<0.05 by Student’s *t* test. Data is representative of three independent experiment (mean ± SD).

**Figure S2.**
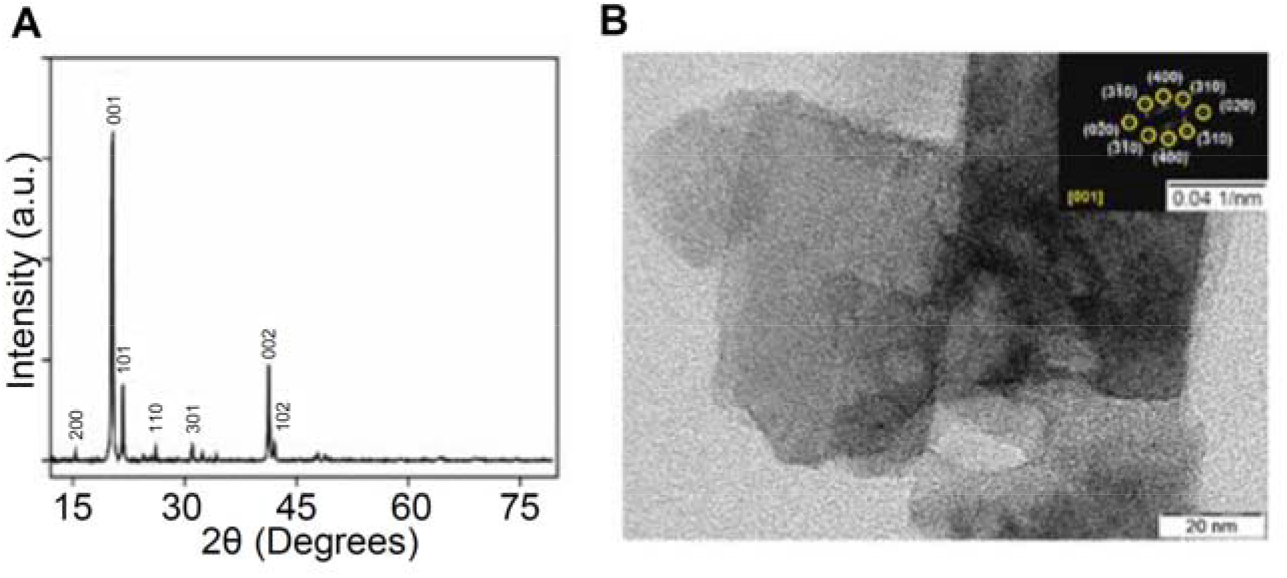
XRD pattern and TEM of Vs. **(A)** Powder XRD of Vs where all the diffraction peaks were indexed to standard V_2_O_5_ orthorhombic phase (JCPDS = 41-1426, space group Pmmn). **(B)** Transmission electron microscopy (TEM) image of Vs thin nanosheets and selected area electron diffraction (SAED) (inset) shows the crystalline nature of the material.

**Figure S3.**
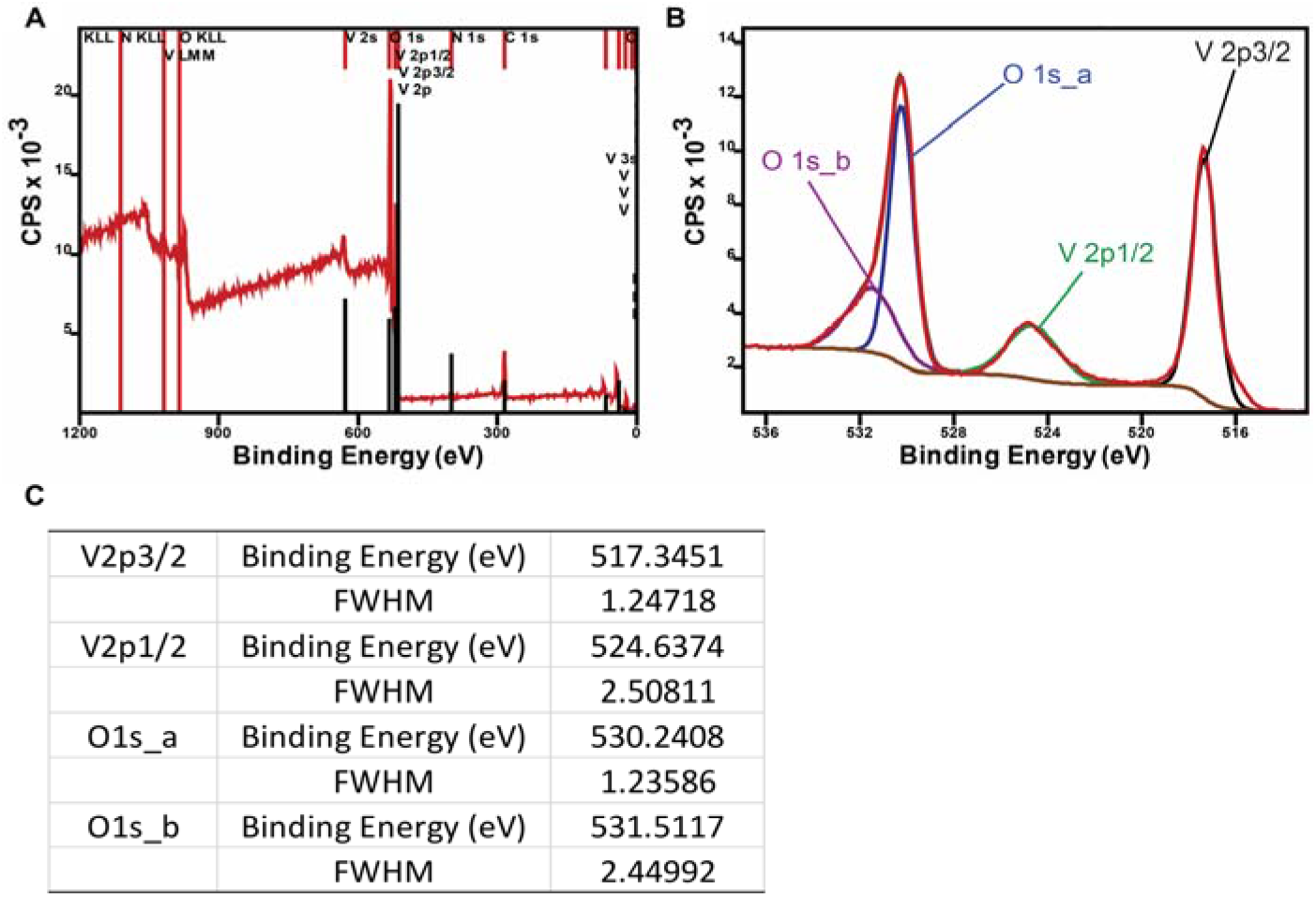
X-Ray Photoelectron Spectroscopy (XPS) of Vs. **(A)** Left: wide spectra and **(B)** right: deconvoluted spectra of oxygen and vanadium peaks. The spectra were calibrated by taking C1s (284.6 eV) as a standard. As described by *Mendialdua* e*t al* (*81, 82*) the difference in binding energies between the O1s core level and the V2p3/2 level (Δ = BE (O1s) – BE (V2p3/2)) was used to determine the oxidation state of V_2_O_5_ nanozymes which confirmed +5 oxidation states of vanadium in Vs. **(C)** Full width half maxima (FWHM) & binding energies of deconvoluted oxygen and vanadium XPS peaks.

**Figure S4.**
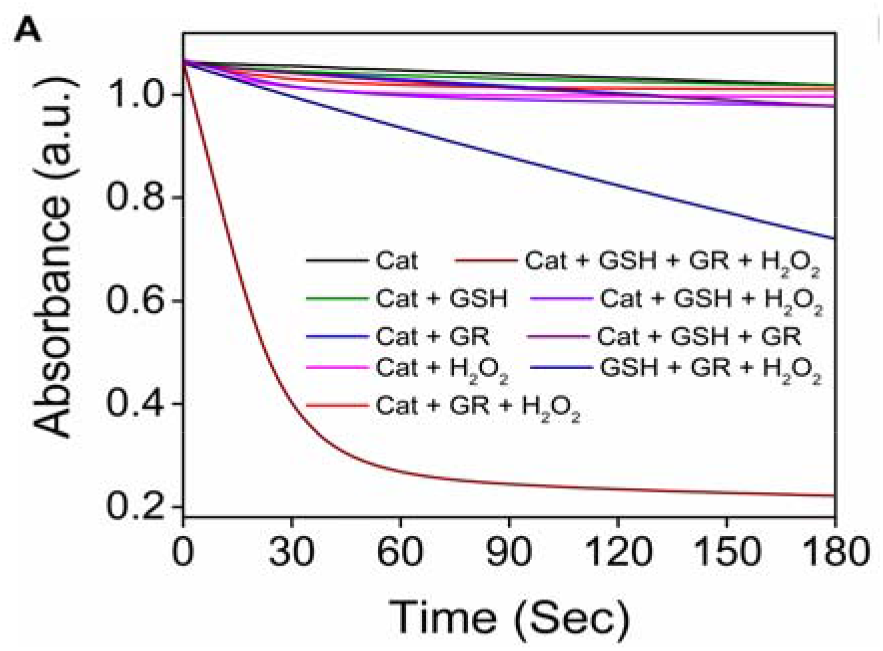
GR- coupled assay for Vs under varied conditions. **(A)** The decrease in absorbance of NADPH (0.2 mM under different assay conditions were monitored during GR coupled assay.

**Figure S5.**
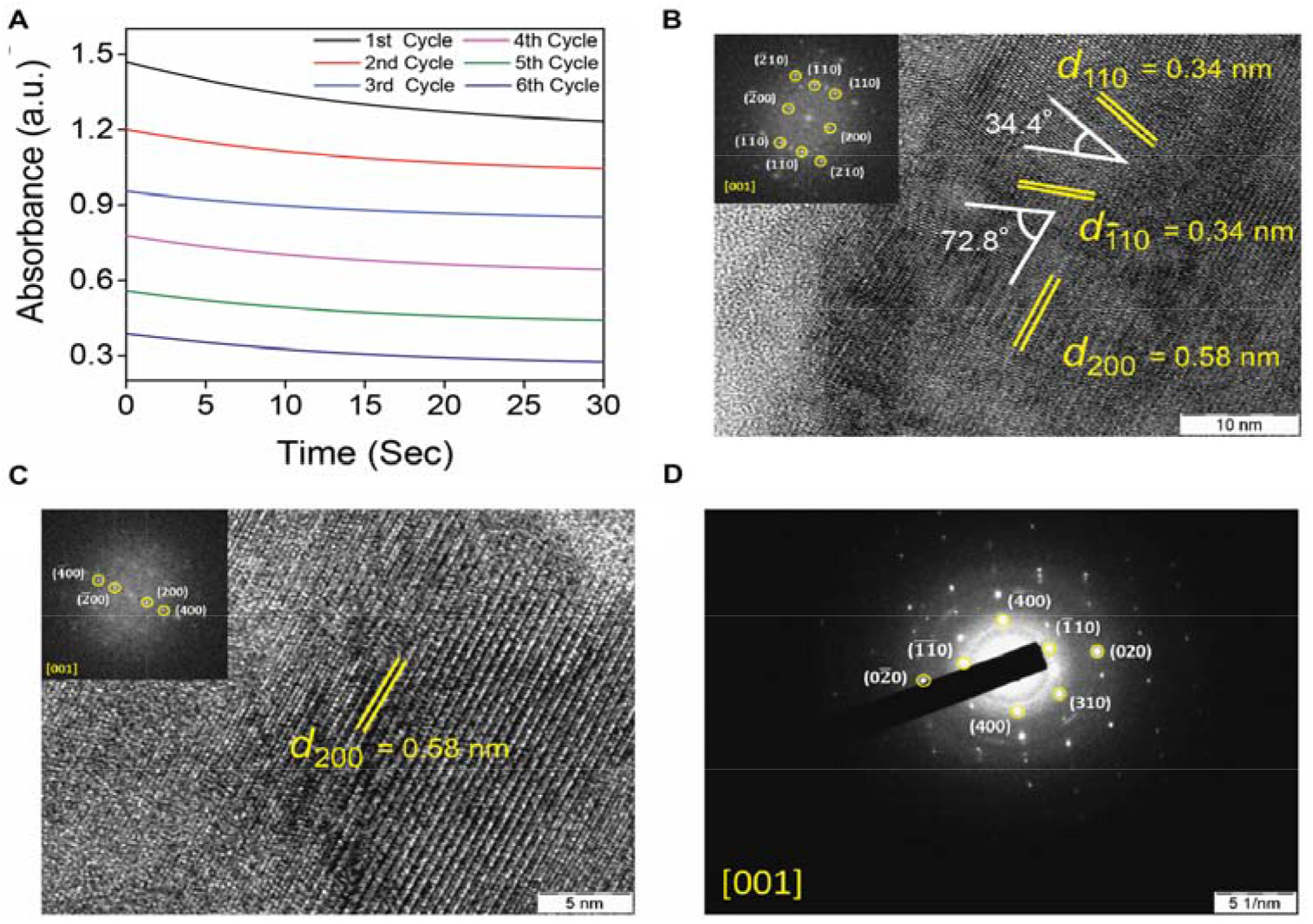
*In vitro* recycling ability of Vs. **(A)** Recycling activity of Vs during multiple rounds of catalysis was analysed by addition of fresh substrates in the reaction mixture up to six cycles. The curves observed due to reduction in NADPH absorbance, for every cycle were parallel to each other, indicating no change in their initial rate and persistence of the activity. This clearly depicts robustness of the catalyst. Conditions used for the assay was sodium phosphate buffer (100 mM, pH 7.4), GSH (2 mM), NADPH (0.2 mM), catalyst (20 ng/μM) at 25°C. High Resolution TEM (HRTEM) image and FFT patterns of Vs before **(B)** and after **(C)** catalysis. **(D)** SAED pattern of Vs after catalysis. The pattern was indexed in a common zone axis [001] which indicates the surface exposed facets are retained after multiple rounds of catalysis.

**Figure S6.**
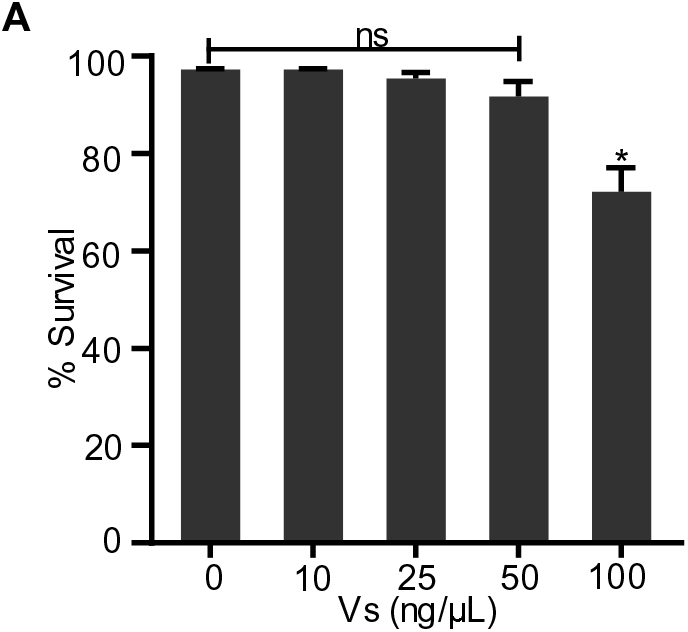
Survival assay for Vs-treated U1 cells. **(A)** U1 cells were treated with increasing concentrations of Vs – 10 to 100 ng/ μL for 15 min and cell survival was analysed flow cytometrically after 24 h by propidium iodide (PI) staining. Experiment is representative of two independent experiments done in triplicate (mean ± SD). *P<0.05, by Mann Whitney test.

**Figure S7.**
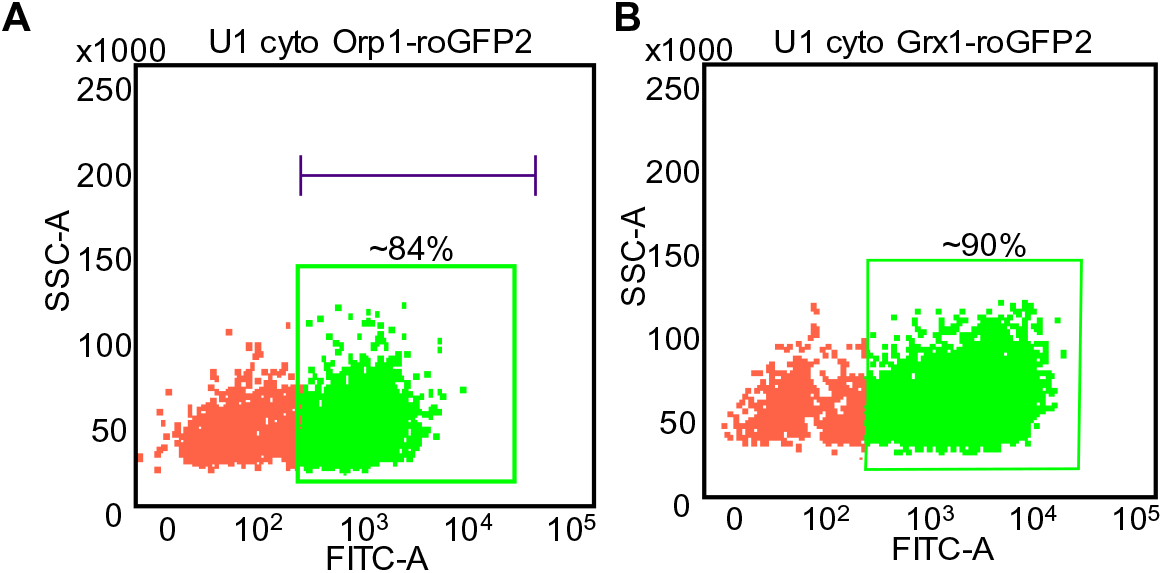
Preparation of Stable cell lines. Dot plot validating the stable expression of the biosensors in **(A)** U1 cyto-Orp1-roGFP2 and **(B)** U1 cyto-Grx1-roGFP2 cell lines.

**Figure S8.**
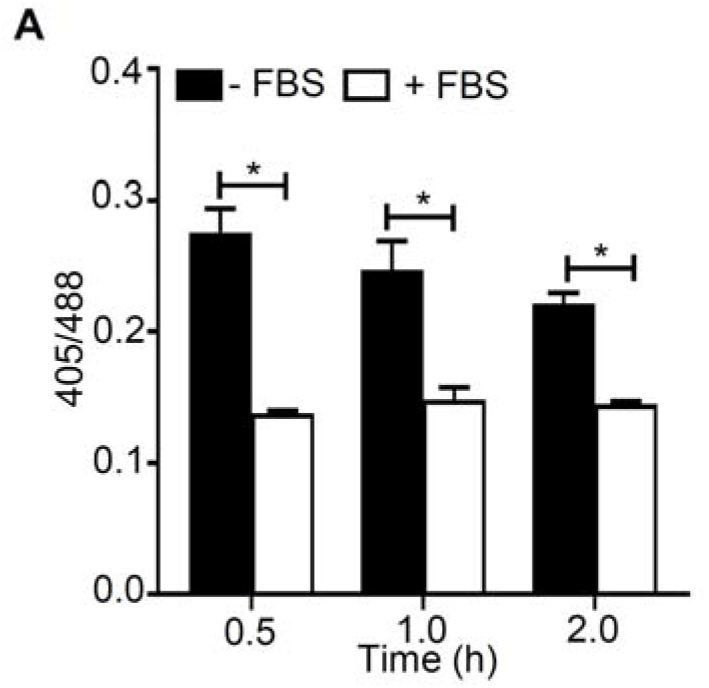
Time dependent induction of Selenium (Se) starvation-mediated oxidative stress. **(A)** U1-Grx1-roGFP2 cells were cultured in serum free medium (to deplete Se) for various time periods and biosensor response was measured by flow cytometry. N=2. Data is representative of two experiments done in duplicates (mean ± SEM). * P<0.05, by Mann Whitney test.

**Figure S9.**
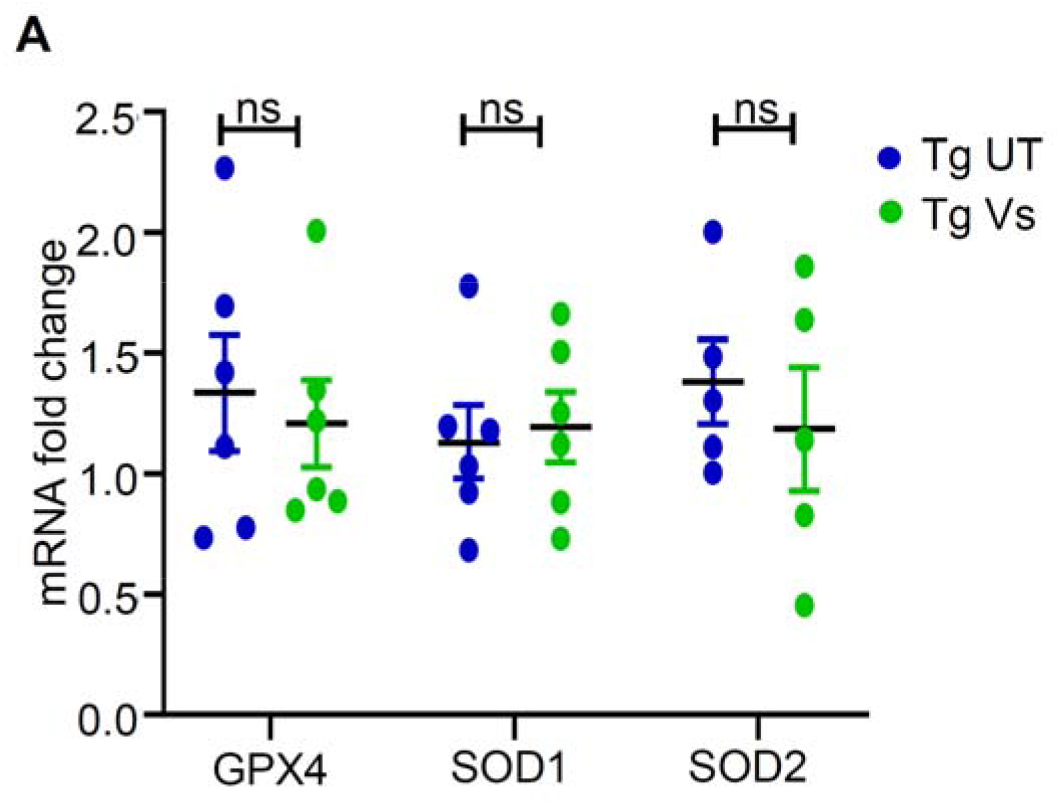
Antioxidant gene expression in lungs of Vs treated Tg26. **(A)** Tg26 mice were left untreated or treated with 1 mg/kg of Vs for 3 h. RNA was isolated from the right lung lobe, reverse transcribed and expression levels of GPX4, SOD1 and SOD2 in Tg UT and Tg Vs animals were analysed. Values were normalised with Tg UT. n=6, ns=non-significant.

### Supplementary Table

**Table S2.**
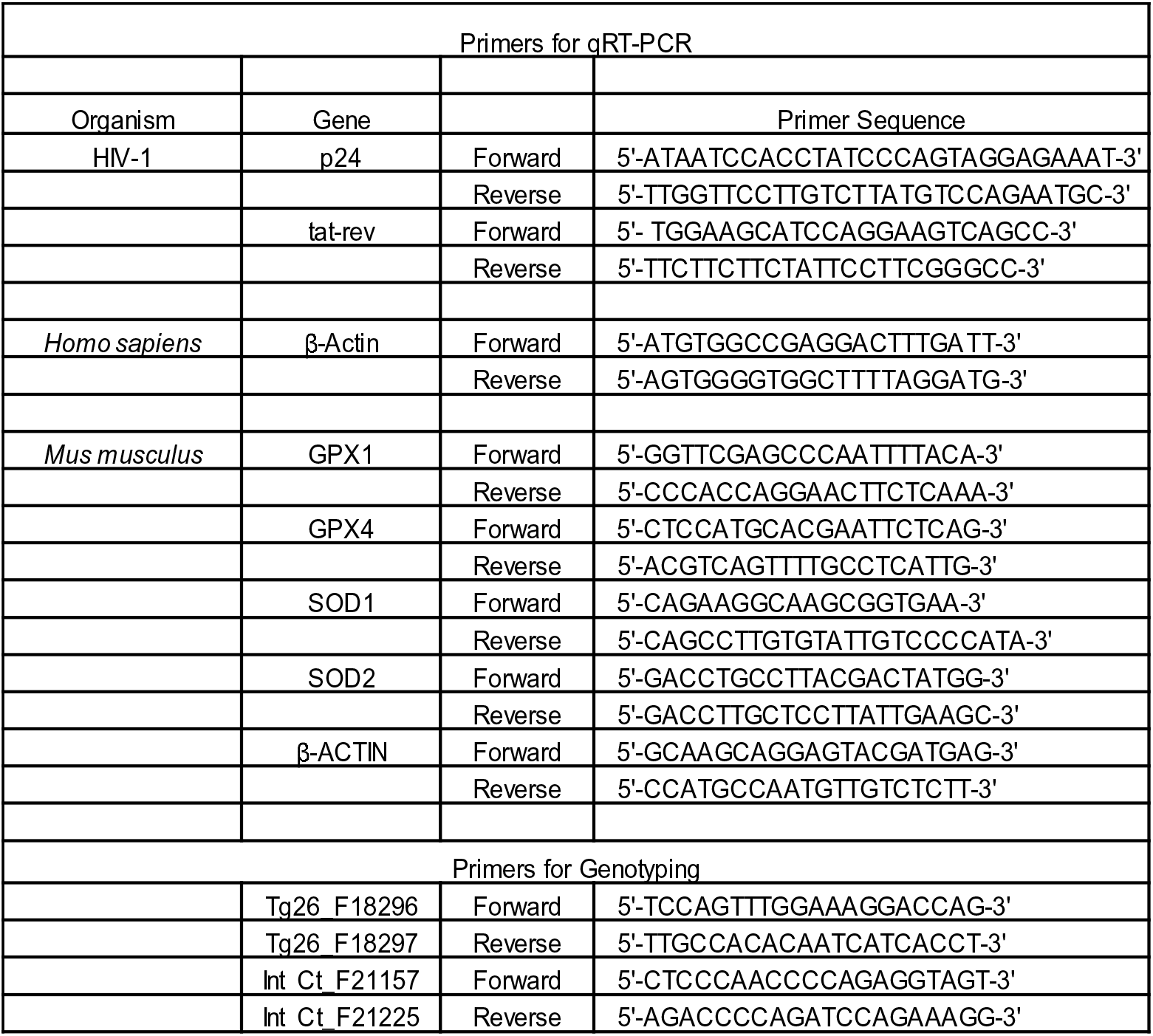
List of primers used in the study.

### Supplementary Materials and Methods

#### Dispersion of V_2_O_5_ NPs

V_2_O_5_ NPs were dispersed in sterile water at a concentration of 2 mg/mL. The dispersion was carried out by sonication using probe sonicator under the following conditions: time – 5 min, amplitude – 5 sec ON, 5 sec OFF. After dispersion, the vanadia NPs form a yellowish colloidal solution.

#### Treatment of cell lines with Vs

Indicated cell lines were treated with different concentrations of Vs at a cell density of 0.2 × 10^6^ cells/mL for 15 min at 37°C in a CO incubator. Following Vs treatment, the cells were washed and re-suspended in complete medium for further culturing or in FACS buffer (1× PBS + 3% FBS) for measuring the antioxidant response by flow cytometry, as required. For experiments with J1.1, Jurkat, or CEM-GFP cell lines, which required long term culturing, Vs treatment was repeated every 24 h, as mentioned above.

#### Internalization of Vs by U1

U1 cells were treated with 50 ng/μL as mentioned above. The cell pellet was re-suspended in 10 mL of dilute nitric acid (HNO_3_; 3 N) and incubated for 16 h to remove cellular organic matter. The samples are analysed by ICP-AES (inductively coupled plasma atomic emission spectroscopy) and their vanadium (V) content was estimated in parts per billion (ppb). The ppb content of vanadium sulphate treated parallelly with HNO_3_ was used as a standard to estimate the cellular V content.

#### Mammalian and bacterial Cell Culture

The human monocytic cell line U937, CD4^+^ T lymphocytic cell line Jurkat (ATCC, Manassas, VA), the chronically infected U1 and J1.1, and CD4^+^ reporter T cell line, CEM-GFP (AIDS Research and Reference Reagent program, NIH, USA) were grown in RPMI 1640 (Cell Clone), with 10% FBS (MP Biomedicals) and 2 mM L-glutamine (MP Biomedicals) supplementation. Vs treatment, transfection, and HIV-1 infection was carried out in Opti MEM media (Hyclone). Differentiation and HIV activation in U1 cells were carried out by treatment with 5 ng/mL of phorbol ester PMA (Sigma) or 1.25 μM prostratin (Sigma). Laboratory strain of *Mtb* H37Rv and clinical isolate JAL2287 (MDR) were cultured in 7H9 (liquid) or 7H11 (solid) synthetic medium supplemented with 0.4% glycerol, 0.1% tween-80 and 10% albumin-dextrose-sodium chloride (ADS) or 0.4% glycerol and 10% oleic acid-albumin-dextrose-catalase (OADC), respectively.

#### Preparation of Stable Cell lines and validation using flow cytometry

Various cell lines stably expressing the cytosolic biosensor Grx1-roGFP2 or Orp1-roGFP2 were prepared by electroporation of 10 ×10^6^ U1 cells with 5 μg of the pMSCVpuro-Grx1-roGFP2 or pMSCVpuro-Orp1-roGFP2 constructs, followed by selection on 350 ng/mL puromycin. The ratiometric responses of the biosensors were measured by excitation at 405 and 488 nm, and recording emission at 510/10 nm, using BD FACSVerse flow cytometer (BD Biosciences). The data were analysed using FACSuite software (BD Biosciences).

#### Assessment of Vs antioxidant activity and redox potential measurement

0.1 ×10^6^ untreated and Vs treated U1-Orp1-roGFP2 or U1-Grx1-roGFP2 cells were exposed to increasing concentrations of H O - 50 and 100 μM or 50, 100 and 200 μM, respectively- and incubated at RT for 2-3 min. These cells were analysed by flow cytometry at excitation of 405 nm (V500) and 488 nm (FITC), while the emission was fixed at 510 nm. Ratio of fluorescence intensities at 405/488 was calculated and normalized using a cell permeable oxidant H_2_O_2_ or the reductant DTT to calculate the responsiveness of both the biosensors.

Intracellular redox potential was measured for cells expressing the Grx1-roGFP2 biosensor, as mentioned earlier (*15*). Briefly, for each experiment, 100% biosensor oxidation or reduction corresponding to maximal and minimal fluorescence intensity ratios was determined by treatment with 10 mM H_2_O_2_ and 10 mM DTT, respectively. The observed ratios were used to determine the degree of biosensor oxidation and ultimately equated in a modified form of the Nernst equation to obtain the intracellular glutathione redox potential (*E_GSH_*).

#### Propidium iodide (PI) staining

U1 cells were treated with increasing concentrations of Vs for 15 min and cultured in complete RPMI medium for 24 h. After 24 h, cells were washed, suspended in 1× PBS, and stained with 3 µM PI for 15 min in the dark. After washing twice with 1× PBS, cells were analyzed on a flow cytometer using the phycoerythrin (PE) detector (575/26 nm) by excitation at 488 nm.

emission.

#### HIV reactivation in U1 cells and qRT-PCR analysis

U1 cells were treated with either 5 ng/mL PMA or 1.25 μM prostratin and incubated at 37°C in a CO incubator. Samples were harvested at 6 h, 12 h, and 24 h post-activation, and RNA was isolated using the Qiagen RNAeasy kit (manufacturer’s protocol). cDNA was synthesized using 400 ng RNA by the Biorad iScript cDNA synthesis kit. qRT-PCR was performed using primers against *gag* transcript (a marker for HIV reactivation). Actin was used as an internal control. To inhibit NF-κB pathway, untreated or Vs-treated cells were exposed to 7.5 µM Bay11-7082 (TCI chemicals) for 12 h.

#### p24 detection by Immunoblotting and ELISA

Untreated or Vs treated HIV-infected Jurkat cells were harvested at 36, 48, and 72 h post infection. Cells were lysed in 300 μL of passive lysis buffer (Promega) supplemented with 1×protease inhibitor cocktail (Roche). Protein was quantified using Pierce™ BCA Protein Assay Kit (Thermo Fisher Scientific). 50 μM lysate (WCL) was mixed with Laemmli buffer, heated at 95°C for 5 min, and was separated on a 12% SDS PAGE gel. Immunoblotting was performed using primary antibodies against HIV-p24 (Abcam; ab9071) and GAPDH (CST; D4C6R) as an internal control. Horse anti-mouse IgG (CST; 7076) was used as the secondary antibody. For ELISA, we collected supernatant from Jurkat cells infected with HIV at 24, 48, and 72 hpi. p24 levels were determined by sandwich ELISA using the J.J. Mitra’s kit as per the manufacturers’ instructions. Standard curve was prepared with known amount p24 and utilised for calculating the viral p24 concentration in the medium.

#### Bacterial Survival assays in U1 cells

For survival experiments of *Mtb* H37Rv and JAL2287 strains in U1 cell lines, the cells were differentiated with 5 ng/mL of PMA for 18 h. Differentiated U1 macrophages were either left untreated or treated with 50 ng/μL of Vs for 15 min, washed, and allowed to rest for 2 h. H37Rv and JAL2287 were opsonized in 50% horse serum for 1 h. Untreated or Vs treated U1 cells were infected at moi 2 for 4 h. After 4 h of incubation, extracellular bacilli were killed by treatment with 0.2 mg/mL amikacin for 1 h. The infected cells were washed three times with 1× PBS and then cultured in complete RPMI media for 24 h. The infected cells were lysed in 0.06% sodium dodecyl sulphate (SDS) and plated on 7H11 plates supplemented with 10% OADC. Bacterial survival was assessed 24 h post infection by enumerating colony forming units (CFUs) on agar media. The initial bacterial burden was determined by plating at 0 h post infection and the percentage increase in bacterial survival was plotted.

#### Isolation and infection of primary CD4^+^ T cells

Peripheral blood mononuclear cells (PBMCs) were isolated using Ficoll-Paque-based density gradient centrifugation from blood samples of healthy donors, donated after informed consent, approved by the IISc Ethical Committee. Primary CD4^+^ T cells were purified from PBMCs using an EasySep human CD4^+^ T cell isolation kit (Stem Cell Technologies, Canada) and cultured for 3 days after isolation at 37°C in 5% CO_2_ in complete media containing RPMI 1640 supplemented with 10% FBS, 100 U/mL interleukin-2 (IL-2) (Peprotech, London, United Kingdom) (specific activity, 10 U/ng), and 1_µg/mL phytohemagglutinin (PHA) (Thermo Fisher Scientific). Subsequently, 250,000 activated primary CD4^+^ T cells, were pre-treated with 25 ng/µL Vs for 15 min and infected with 0.1 moi pNL4.3 virus by spinoculation at 1000 g for 90 min at 32°C. Cells were then washed and replenished with complete media containing 100 U/mL IL-2. Vs treatment was repeated every 48 h. To quantify the virion release, supernatant was harvested from infected cells and centrifuged at 400 g for 10 min and virus concentration was estimated by HIV-1 p24 ELISA.

#### Animal experiments

Heterozygous HIV transgenic mouse Tg26 containing HIV proviral DNA with mutated 3 kb region of *gag* and *pol* genes was used (*64*). In order to create a more stable line, the HIV Tg26 (FVB/N) was backcrossed with C57BL/6 for eight generations to create the current Tg26 by Dr. Roy L. Sutliff (Veterans Affairs Medical Center, Emory University, Atlanta, GA). HIV Tg26 breeding colony was maintained in the BSL3 animal facility at CIDR, IISc, according to the guidelines set by Institutional Animal Ethical Committee. Genotyping was done at 4 weeks of age by tail vein PCR. 6-9 weeks old sex-matched transgenic mice were used for the experiment. Wild type (*Wt*) littermates were used as controls. 1 mg/kg body weight Vs was administered by intra-peritoneal injection in Tg26 treatment group. Control and treated animals were euthanized at 3 h post Vs treatment.

#### Pulmonary edema analysis

Left lobe of the lungs from 8 mice in each group-untreated C57BL/6 Tg26 and Vs treated Tg26 were removed aseptically and weighed in 1.5 mL microfuge tubes. The lungs were left for 16-24 h in a dry oven at 55°C for desiccation and the dry weight of the tissue was recorded. The ratio of lung wet weight to dry weight was plotted for each group.

#### ROS staining of lung homogenates

Untreated WT, untreated Tg26, and Vs treated Tg26 mice were euthanized at 3 h post Vs treatment and a portion of the right lobe of the lung was aseptically removed. Single cell suspensions for lung homogenates were prepared for DCFDA staining and flow cytometry analysis as per an earlier study (*73*). Briefly, the tissue was minced and digested in serum free RPMI containing 0.2 mg/mL Liberase DL and 0.1 mg/mL DNase I for 60 min at 37°C and agitation at 180 rpm. The minced tissue was mechanically disrupted using GentleMACS tissue dissociator to obtain a finer suspension. Larger cell clumps were removed by passing the suspension through a 40 μm cell strainer (BD Falcon). Red blood cells were removed by incubating the cells in RBC lysis buffer.

#### qRT-PCR analysis from animal tissues

RNA was isolated from a part of the right lobe of the lungs using Trizol reagent (Ambion). 1 μg RNA was reverse transcribed using the Biorad iScript cDNA synthesis kit. qRT-PCR was performed using primers against GPX1, GPX4, CAT, SOD1, and SOD2 to assess the host antioxidant response under different treatment groups. β-Actin was used as an internal control.

## References

1. H. Wei, E. Wang, Nanomaterials with enzyme-like characteristics (nanozymes): next-generation artificial enzymes. Chem. Soc. Rev.42, 6060–6093 (2013).

2. R. de la Rica, M. M. Stevens, Plasmonic ELISA for the ultrasensitive detection of disease biomarkers with the naked eye. Nat. nanotechno. 7, 821–824 (2012)

3. O. Salata, Applications of nanoparticles in biology and medicine. J. Nanobiotechnology 2, 3 (2004).

4. C. Korsvik, S. Patil, S. Seal, W. T. Self, Superoxide dismutase mimetic properties exhibited by vacancy engineered ceria nanoparticles. Chem. Commun. (Camb), 1056–1058 (2007).

5. M. Das, S. Patil, N. Bhargava, J. F. Kang, L. M. Riedel, S. Seal, J. J. Hickman, Auto-catalytic ceria nanoparticles offer neuroprotection to adult rat spinal cord neurons. Biomaterials 28, 1918–1925 (2007).

6. L. Gao, Y. Liu, D. Kim, Y. Li, G. Hwang, P. C. Naha, D. P. Cormode, H. Koo, Nanocatalysts promote Streptococcus mutans biofilm matrix degradation and enhance bacterial killing to suppress dental caries in vivo. Biomaterials 101, 272–284 (2016).

7. S. Zanganeh, G. Hutter, R. Spitler, O. Lenkov, M. Mahmoudi, A. Shaw, J. S. Pajarinen, H. Nejadnik, S. Goodman, M. Moseley, L. M. Coussens, H. E. Daldrup-Link, Iron oxide nanoparticles inhibit tumour growth by inducing pro-inflammatory macrophage polarization in tumour tissues. Nat. Nanotechnol. 11, 986–994 (2016).

8. A. A. Vernekar, D. Sinha, S. Srivastava, P. U. Paramasivam, P. D’Silva, G. Mugesh, An antioxidant nanozyme that uncovers the cytoprotective potential of vanadia nanowires. Nat. Commun. 5, 5301 (2014).

9. A. Perl, K. Banki, Genetic and metabolic control of the mitochondrial transmembrane potential and reactive oxygen intermediate production in HIV disease. Antioxid. Redox Signal. 2, 551–573 (2000).

10. F. J. Staal, M. Roederer, L. A. Herzenberg, L. A. Herzenberg, Intracellular thiols regulate activation of nuclear factor kappa B and transcription of human immunodeficiency virus. Proc. Natl. Acad. Sci.U.S.A. 87, 9943–9947 (1990).

11. C. W. Pyo, Y. L. Yang, N. K. Yoo, S. Y. Choi, Reactive oxygen species activate HIV long terminal repeat via post-translational control of NF-kappaB. Biochem. Biophys. Res. Commun. 376, 180–185 (2008).

12. H. P. Eck, H. Gmunder, M. Hartmann, D. Petzoldt, V. Daniel, W. Droge, Low concentrations of acid-soluble thiol (cysteine) in the blood plasma of HIV-1-infected patients. Biol. Chem. Hoppe Seyler 370, 101–108 (1989).

13. R. Buhl, H. A. Jaffe, K. J. Holroyd, F. B. Wells, A. Mastrangeli, C. Saltini, A. M. Cantin, R. G. Crystal, Systemic glutathione deficiency in symptom-free HIV-seropositive individuals. Lancet 2, 1294–1298 (1989).

14. L. A. Herzenberg, S. C. De Rosa, J. G. Dubs, M. Roederer, M. T. Anderson, S. W. Ela, S. C. Deresinski, L. A. Herzenberg, Glutathione deficiency is associated with impaired survival in HIV disease. Proc. Natl. Acad. Sci. U.S.A. 94, 1967–1972 (1997).

15. A. Bhaskar, M. Munshi, S. Z. Khan, S. Fatima, R. Arya, S. Jameel, A. Singh, Measuring glutathione redox potential of HIV-1-infected macrophages. J. Biol. Chem. 290, 1020–1038 (2015).

16. M. P. Look, J. K. Rockstroh, G. S. Rao, K. A. Kreuzer, S. Barton, H. Lemoch, T. Sudhop, J. Hoch, K. Stockinger, U. Spengler, T. Sauerbruch, Serum selenium, plasma glutathione (GSH) and erythrocyte glutathione peroxidase (GSH-Px)-levels in asymptomatic versus symptomatic human immunodeficiency virus-1 (HIV-1)-infection. Eur. J. Clin. Nutr. 51, 266–272 (1997).

17. L. Zhao, A. G. Cox, J. A. Ruzicka, A. A. Bhat, W. Zhang, E. W. Taylor, Molecular modeling and in vitro activity of an HIV-1-encoded glutathione peroxidase. Proc. Natl. Acad. Sci. U.S.A. 97, 6356–6361 (2000).

18. I. Cohen, P. Boya, L. Zhao, D. Metivier, K. Andreau, J. L. Perfettini, J. G. Weaver, A. Badley, E. W. Taylor, G. Kroemer, Anti-apoptotic activity of the glutathione peroxidase homologue encoded by HIV-1. Apoptosis 9, 181–192 (2004).

19. S. C. De Rosa, M. D. Zaretsky, J. G. Dubs, M. Roederer, M. Anderson, A. Green, D. Mitra, N. Watanabe, H. Nakamura, I. Tjioe, S. C. Deresinski, W. A. Moore, S. W. Ela, D. Parks, L. A. Herzenberg, L. A. Herzenberg, N-acetylcysteine replenishes glutathione in HIV infection. Eur. J. Clin. Invest. 30, 915–929 (2000).

20. A. Witschi, E. Junker, C. Schranz, R. F. Speck, B. H. Lauterburg, Supplementation of N-acetylcysteine fails to increase glutathione in lymphocytes and plasma of patients with AIDS. AIDS Res. Hum. Retroviruses 11, 141–143 (1995).

21. C. Sappey, S. Legrand-Poels, M. Best-Belpomme, A. Favier, B. Rentier, J. Piette, Stimulation of glutathione peroxidase activity decreases HIV type 1 activation after oxidative stress. AIDS Res. Hum. Retroviruses 10, 1451–1461 (1994).

22. P. A. Sandstrom, J. Murray, T. M. Folks, A. M. Diamond, Antioxidant defenses influence HIV-1 replication and associated cytopathic effects. Free Radic. Biol. Med. 24, 1485–1491 (1998).

23. C. Cao, Y. Leng, W. Huang, X. Liu, D. Kufe, Glutathione peroxidase 1 is regulated by the c-Abl and Arg tyrosine kinases. J. Biol. Chem. 278, 39609–39614 (2003).

24. T. Wiedenmann, N. Dietrich, T. Fleming, S. Altamura, L. E. Deelman, R. H. Henning, M. U. Muckenthaler, P. P. Nawroth, H. P. Hammes, A. H. Wagner, M. Hecker, Modulation of glutathione peroxidase activity by age-dependent carbonylation in glomeruli of diabetic mice. J. Diabetes Complications 32, 130–138 (2018).

25. W. H. Yang, S. Y. Park, S. Ji, J. G. Kang, J. E. Kim, H. Song, I. Mook-Jung, K. Choe, J. W. Cho, O-GlcNAcylation regulates hyperglycemia-induced GPX1 activation. Biochem. Biophys. Res. Commun. 391, 756–761 (2010).

26. S. Ghosh, P. Roy, N. Karmodak, E. D. Jemmis, G. Mugesh, Nanoisozymes: Crystal-Facet-Dependent Enzyme-Mimetic Activity of V_2_O_5_ Nanomaterials. Angewandte Chemie 57, 4510–4515 (2018).

27. J. R. Morones, J. L. Elechiguerra, A. Camacho, K. Holt, J. B. Kouri, J. T. Ramirez, M. J. Yacaman, The bactericidal effect of silver nanoparticles. Nanotechnology 16, 2346–2353 (2005).

28. W. Jiang, B. Y. Kim, J. T. Rutka, W. C. Chan, Nanoparticle-mediated cellular response is size-dependent. Nat. Nanotechnol. 3, 145–150 (2008).

29. Y. Pan, S. Neuss, A. Leifert, M. Fischler, F. Wen, U. Simon, G. Schmid, W. Brandau, W. Jahnen-Dechent, Size-dependent cytotoxicity of gold nanoparticles. Small 3, 1941–1949 (2007).

30. M. Auffan, J. Rose, J. Y. Bottero, G. V. Lowry, J. P. Jolivet, M. R. Wiesner, Towards a definition of inorganic nanoparticles from an environmental, health and safety perspective. Nat. Nanotechnol. 4, 634–641 (2009).

31. X. Zhou, W. Xu, G. Liu, D. Panda, P. Chen, Size-dependent catalytic activity and dynamics of gold nanoparticles at the single-molecule level. J. Am. Chem. Soc. 132, 138–146 (2010).

32. B. Zhou, D, He, Raman spectrum of vanadium pentoxide from density functional perturbation theory. J. Raman Spectrosc. 39, 1475–1481 (2008).

33. W. Avansi, J. L. Q. Maia, C. Ribeiro, E. R. Leite, V. R. Mastelaro, Local structure study of vanadium pentoxide 1D-nanostructures. J. Nanopart. Res. 13, (2011).

34. C. Sanchez, J. Livage, G. Lucazeau, Infrared and Raman study of amorphous V_2_O_5_. J. Raman Spectrosc. 12, 68–72 (1982).

35. T. M. Folks, J. Justement, A. Kinter, C. A. Dinarello, A. S. Fauci, Cytokine-induced expression of HIV-1 in a chronically infected promonocyte cell line. Science 238, 800–802 (1987).

36. G. Poli, A. Kinter, J. S. Justement, J. H. Kehrl, P. Bressler, S. Stanley, A. S. Fauci, Tumor necrosis factor alpha functions in an autocrine manner in the induction of human immunodeficiency virus expression. Proc. Natl. Acad. Sci. U.S.A. 87, 782–785 (1990).

37. M. Gutscher, A. L. Pauleau, L. Marty, T. Brach, G. H. Wabnitz, Y. Samstag, A. J. Meyer, T. P. Dick, Real-time imaging of the intracellular glutathione redox potential. Nat. Methods 5, 553–559 (2008).

38. B. Morgan, M. C. Sobotta, T. P. Dick, Measuring E(GSH) and H_2_O_2_ with roGFP2-based redox probes. Free Radic. Biol. Med. 51, 1943–1951 (2011).

39. M. Gutscher, M. C. Sobotta, G. H. Wabnitz, S. Ballikaya, A. J. Meyer, Y. Samstag, T. P. Dick, Proximity-based protein thiol oxidation by H_2_O_2_-scavenging peroxidases. J. Biol. Chem. 284, 31532–31540 (2009).

40. R. Brigelius-Flohe, M. Maiorino, Glutathione peroxidases Biochim. Biophys. Acta 1830, 3289–3303 (2013).

41. R. Drew, J. O. Miners, The effects of buthionine sulphoximine (BSO) on glutathione depletion and xenobiotic biotransformation. Biochemical Pharmacology 33, 2989–2994 (1984).

42. S. Legrand-Poels, D. Vaira, J. Pincemail, A. van de Vorst, J. Piette, Activation of human immunodeficiency virus type 1 by oxidative stress. AIDS Res. Hum. Retroviruses 6, 1389–1397 (1990).

43. C. H. Kim, S. Gollapudi, A. Kim, T. Lee, S. Gupta, Role of protein kinase C-beta isozyme in activation of latent human immunodeficiency virus type 1 in promonocytic U1 cells by phorbol-12-myristate acetate. AIDS Res. Hum. Retroviruses 12, 1361–1366 (1996).

44. R. J. Gulakowski, J. B. McMahon, R. W. Buckheit, Jr., K. R. Gustafson, M. R. Boyd, Antireplicative and anticytopathic activities of prostratin, a non-tumor-promoting phorbol ester, against human immunodeficiency virus (HIV). Antiviral Res. 33, 87–97 (1997).

45. A. Campa, G. Shor-Posner, F. Indacochea, G. Zhang, H. Lai, D. Asthana, G. B. Scott, M. K. Baum, Mortality risk in selenium-deficient HIV-positive children. J. Acquir. Immune Defic. Syndr. Hum. Retrovirol. 20, 508–513 (1999).

46. M. P. Look, J. K. Rockstroh, G. S. Rao, K. A. Kreuzer, U. Spengler, T. Sauerbruch, Serum selenium versus lymphocyte subsets and markers of disease progression and inflammatory response in human immunodeficiency virus-1 infection. Biol. Trace Elem. Res. 56, 31–41 (1997).

47. A. Gervaix, D. West, L. M. Leoni, D. D. Richman, F. Wong-Staal, J. Corbeil, A new reporter cell line to monitor HIV infection and drug susceptibility in vitro. Proc. Natl. Acad. Sci. U.S.A. 94, 4653–4658 (1997).

48. P. Tyagi, V. K. Pal, R. Agrawal, S. Singh, S. Srinivasan, A. Singh, *Mycobacterium tuberculosis* Reactivates HIV-1 via Exosome-Mediated Resetting of Cellular Redox Potential and Bioenergetics. mBio 11, (2020).

49. E. P. Amaral, E. L. Conceicao, D. L. Costa, M. S. Rocha, J. M. Marinho, M. Cordeiro-Santos, M. R. D’Imperio-Lima, T. Barbosa, A. Sher, B. B. Andrade, N-acetyl-cysteine exhibits potent anti-mycobacterial activity in addition to its known anti-oxidative functions. BMC Microbiol. 16, 251 (2016).

50. G. Teskey, R. Cao, H. Islamoglu, A. Medina, C. Prasad, R. Prasad, A. Sathananthan, M. Fraix, S. Subbian, L. Zhong, V. Venketaraman, The Synergistic Effects of the Glutathione Precursor, NAC and First-Line Antibiotics in the Granulomatous Response Against *Mycobacterium tuberculosis*. Front. Immunol. 9, 2069 (2018).

51. D. Kumar, L. Nath, M. A. Kamal, A. Varshney, A. Jain, S. Singh, K. V. Rao, Genome-wide analysis of the host intracellular network that regulates survival of *Mycobacterium tuberculosis*. Cell 140, 731–743 (2010).

52. M. M. Kulkarni, Digital multiplexed gene expression analysis using the NanoString nCounter system. Curr. Protoc. Mol. Biol. Chapter 25, Unit25B 10 (2011).

53. R. Eid, E. Boucher, N. Gharib, C. Khoury, N. T. Arab, A. Murray, P. G. Young, C. A. Mandato, M. T. Greenwood, Identification of human ferritin, heavy polypeptide 1 (FTH1) and yeast RGI1 (YER067W) as pro-survival sequences that counteract the effects of Bax and copper in *Saccharomyces cerevisiae*. Exp. Cell. Res. 342, 52–61 (2016).

54. H. Sato, A. Shiiya, M. Kimata, K. Maebara, M. Tamba, Y. Sakakura, N. Makino, F. Sugiyama, K. Yagami, T. Moriguchi, S. Takahashi, S. Bannai, Redox imbalance in cystine/glutamate transporter-deficient mice. J. Biol. Chem. 280, 37423–37429 (2005).

55. C. Espinosa-Diez, V. Miguel, D. Mennerich, T. Kietzmann, P. Sanchez-Perez, S. Cadenas, S. Lamas, Antioxidant responses and cellular adjustments to oxidative stress. Redox biology 6, 183–197 (2015).

56. K. A. Roebuck, D. S. Gu, M. F. Kagnoff, Activating protein-1 cooperates with phorbol ester activation signals to increase HIV-1 expression. AIDS 10, 819–826 (1996).

57. A. J. Henderson, K. L. Calame, CCAAT/enhancer binding protein (C/EBP) sites are required for HIV-1 replication in primary macrophages but not CD4(+) T cells. Proc. Natl. Acad. Sci. U.S.A. 94, 8714–8719 (1997).

58. R. Hu, N. Oyaizu, S. Than, V. S. Kalyanaraman, X. P. Wang, S. Pahwa, HIV-1 gp160 induces transforming growth factor-beta production in human PBMC. Clin. Immunol. Immunopathol. 80, 283–289 (1996).

59. G. Herbein, U. Mahlknecht, F. Batliwalla, P. Gregersen, T. Pappas, J. Butler, W. A. O’Brien, E. Verdin, Apoptosis of CD8+ T cells is mediated by macrophages through interaction of HIV gp120 with chemokine receptor CXCR4. Nature 395, 189–194 (1998).

60. C. Amiel, E. Darcissac, M. J. Truong, J. Dewulf, M. Loyens, Y. Mouton, A. Capron, G. M. Bahr, Interleukin-16 (IL-16) inhibits human immunodeficiency virus replication in cells from infected subjects, and serum IL-16 levels drop with disease progression. J. Infect. Dis. 179, 83–91 (1999)

61. W. Choe, D. J. Volsky, M. J. Potash, Induction of rapid and extensive beta-chemokine synthesis in macrophages by human immunodeficiency virus type 1 and gp120, independently of their coreceptor phenotype. J. Virol. 75, 10738–10745 (2001)

62. S. Z. Khan, N. Hand, S. L. Zeichner, Apoptosis-induced activation of HIV-1 in latently infected cell lines. Retrovirology 12, 42 (2015).

63. K. Devadas, N. J. Hardegen, L. M. Wahl, I. K. Hewlett, K. A. Clouse, K. M. Yamada, S. Dhawan, Mechanisms for Macrophage-Mediated HIV-1 Induction. J. Immunol. 173 (11), 6735–6744 (2004).

64. P. Dickie, J. Felser, M. Eckhaus, J. Bryant, J. Silver, N. Marinos, A. L. Notkins, HIV-associated nephropathy in transgenic mice expressing HIV-1 genes. Virology 185, 109–119 (1991).

65. B. A. Jacob, K. M. Porter, S. C. Elms, P. Y. Cheng, D. P. Jones, R. L. Sutliff, HIV-1-induced pulmonary oxidative and nitrosative stress: exacerbated response to endotoxin administration in HIV-1 transgenic mouse model. Am. J. Physiol. Lung. Cell. Mol. Physiol. 291, L811–819 (2006).

66. J. B. Kopp, M. E. Klotman, S. H. Adler, L. A. Bruggeman, P. Dickie, N. J. Marinos, M. Eckhaus, J. L. Bryant, A. L. Notkins, P. E. Klotman, Progressive glomerulosclerosis and enhanced renal accumulation of basement membrane components in mice transgenic for human immunodeficiency virus type 1 genes. Proc. Natl. Acad. Sci. U.S.A. 89, 1577–1581 (1992).

67. S. G. Deeks, R. Tracy, D. C. Douek, Systemic effects of inflammation on health during chronic HIV infection. Immunity 39, 633–645 (2013).

68. S. Ghosh, Prasad, S., Mugesh, G., Understanding the role of oxo and peroxido species in the glutathione peroxidase (GPx)-like activity of metal based nanozymes. Inorganica Chimica Acta 484, 283–290 (2019).

69. V. N. Gladyshev, T. C. Stadtman, D. L. Hatfield, K. T. Jeang, Levels of major selenoproteins in T cells decrease during HIV infection and low molecular mass selenium compounds increase. Proc. Natl. Acad. Sci. U.S.A. 96, 835–839 (1999).

70. J. Q. Wu, T. R. Sasse, G. Wolkenstein, V. Conceicao, M. M. Saksena, M. Soedjono, S. S. Perera, B. Wang, D. E. Dwyer, N. K. Saksena, Transcriptome analysis of primary monocytes shows global down-regulation of genetic networks in HIV viremic patients versus long-term non-progressors. Virology 435, 308–319 (2013)

71. S. Y. Lee, Y. K. Park, C. H. Yoon, K. Kim, K. C. Kim, Meta-analysis of gene expression profiles in long-term non-progressors infected with HIV-1. BMC Med. Genomics 12, 3 (2019)

72. L. L. Zhang, Z. N. Zhang, X. Wu, Y. J. Jiang, Y. J. Fu, H. Shang, Transcriptomic meta-analysis identifies gene expression characteristics in various samples of HIV-infected patients with nonprogressive disease. J. Transl. Med. 15, 191 (2017).

73. R. Mishra, S. Kohli, N. Malhotra, P. Bandyopadhyay, M. Mehta, M. Munshi, V. Adiga, V. K. Ahuja, R. K. Shandil, R. S. Rajmani, A. S. N. Seshasayee, A. Singh, Targeting redox heterogeneity to counteract drug tolerance in replicating *Mycobacterium tuberculosis*. Sci. Transl. Med. 11, (2019).

74. E. P. Amaral, D. L. Costa, S. Namasivayam, N. Riteau, O. Kamenyeva, L. Mittereder, K. D. Mayer-Barber, B. B. Andrade, A. Sher, A major role for ferroptosis in *Mycobacterium tuberculosis*-induced cell death and tissue necrosis. J. Exp. Med. 216, 556–570 (2019).

75. L. Xu, Y. Liu, Z. Chen, W. Li, Y. Liu, L. Wang, Y. Liu, X. Wu, Y. Ji, Y. Zhao, L. Ma, Y. Shao, C. Chen, Surface-engineered gold nanorods: promising DNA vaccine adjuvant for HIV-1 treatment. Nano letters 12, 2003–2012 (2012).

76. M. C. Bowman, T. E. Ballard, C. J. Ackerson, D. L. Feldheim, D. M. Margolis, C. Melander, Inhibition of HIV fusion with multivalent gold nanoparticles. J Am. Chem. Soc. 130, 6896–6897 (2008).

77. H. H. Lara, N. V. Ayala-Nunez, L. Ixtepan-Turrent, C. Rodriguez-Padilla, Mode of antiviral action of silver nanoparticles against HIV-1. J. nanobiotechnology 8, 1 (2010).

78. H. H. Lara, N. V Ayala-Núñez, L. C. I. Turrent, C. R. Padilla, Bactericidal effect of silver nanoparticles against multidrug-resistant bacteria. World Journal of Microbiology and Biotechnology 26, 615–621 (2010).

79. Mandas A, Iorio EL, Congiu MG, Balestrieri C, Mereu A, Cau D, Dessì S, Curreli N. Oxidative imbalance in HIV-1 infected patients treated with antiretroviral therapy. J Biomed Biotechnol. (2009).

80. M. M. Elsheikh, Y. Tang, D. Li, G. Jiang, Deep latency: A new insight into a functional HIV cure. EBioMedicine 45, 624–629 (2019).

81. J. Mendialdua, R. Casanova, Y. Barbaux, XPS studies of V_2_O_5_, V_6_O_13_, VO_2_ and V_2_O_3_. J. Electron Spectrosc. Relat. Phenom. 71, 249–261 (1995).

82. E. Hryha, E. Rutqvist, L. Nyborg, Stoichiometric vanadium oxides studied by XPS. Surf. Interface Anal. 441022–1025 (2012).

